# Asparagine endopeptidase promotes radioresistance in breast cancer through ATR pathway modulation

**DOI:** 10.1101/2025.01.13.632704

**Authors:** Macarena Morillo-Huesca, Mercedes Tomé, Colin Watts, Pablo Huertas, Gema Moreno-Bueno, Raúl V. Durán, Jonathan Martinez-Fabregas

## Abstract

**Background:** Tumor resistance represents a major challenge in the current oncology landscape. Asparagine endopeptidase (AEP) overexpression correlates with worse prognosis and reduced overall survival in most human solid tumors. However, the underlying mechanisms of the connection between AEP and reduced overall survival in cancer patients remain unclear.

**Methods:** High-throughput proteomics, cellular and molecular biology approaches and clinical data from breast cancer (BC) patients were used to identify novel, biologically relevant AEP targets. Immunoblotting and qPCR analyses were used to quantify mRNA and protein levels. Flow cytometry, confocal microscopy, chemical inhibitors, siRNA- and shRNA-silencing and DNA repair assays were used as functional assays. In-silico analyses using the TCGA BC dataset and immunofluorescence assays in an independent cohort of invasive ductal (ID) BC patients were used to validate the clinical relevance of our findings.

**Results:** Here we showed a dual role for AEP in genomic stability and radiotherapy resistance in BC patients by suppressing ATR and PPP1R10 levels. Reduced ATR and PPP1R10 levels were found in BC patients expressing high AEP levels and correlated with worst prognosis. Mechanistically, AEP suppresses ATR levels, reducing DNA damage-induced cell death, and PPP1R10 levels, promoting Chek1/P53 cell cycle checkpoint activation, allowing BC cells to efficiently repair DNA. Functional studies revealed AEP-deficiency results in genomic instability, increased DNA damage signaling, reduced Chek1/P53 activation, impaired DNA repair and cell death, with phosphatase inhibitors restoring the DNA damage response in AEP-deficient BC cells. Furthermore, AEP inhibition sensitized BC cells to the chemotherapeutic reagents cisplatin and etoposide. Immunofluorescence assays in an independent cohort of IDBC patients showed increased AEP levels in ductal cells. These analyses showed that higher AEP levels in radioresistant IDBC patients result in ATR nuclear eviction, revealing AEP^high^/ATR^low^ protein levels as an efficient predictive biomarker for the stratification of radioresistant patients.

**Conclusion:** The newly identified AEP/ATR/PPP1R10 axis plays a dual role in genomic stability and radiotherapy resistance in BC. Our work provides new clues to the underlying mechanisms of tumor resistance and strong evidence validating the AEP/ATR axis as a novel predictive biomarker and therapeutic target for the stratification and treatment of radioresistant BC patients.

## Background

Despite the contribution of technical advances in both chemotherapy and radiotherapy strategies to improvements in treatment outcomes and the quality of life of cancer patients [1], tumor resistance remains a significant challenge in the current oncology landscape. This is because 80-90% of deaths in cancer patients are associated to cancer resistance [2, 3], a complex phenomenon linked to a plethora of cellular alterations, including increased DNA repair, defects in the induction of apoptosis, and autophagy [1, 4]. Therefore, the identification of novel druggable players involved in resistance could allow us to improve cancer therapies by sensitizing cancer cells to current treatments.

Proteases are associated with tumor progression across cancer types. Although, initially considered to promote tumor invasion through extracellular matrix degradation, emerging evidence has revealed unexpected functions for these proteases during the onset and progression of cancer [5]. Unfortunately, the use of broad-range protease inhibitors in cancer therapy has failed to improve patient outcomes [6, 7]. Therefore, the identification of the relevant proteases associated with specific cancer types and the characterization of their molecular targets are paramount for their translation to the clinical setting. Among proteases, AEP is the only known protease that specifically hydrolyzes asparaginyl peptide bonds, and to a lesser extent, aspartyl peptide bonds [8], thus suggesting that AEP has specific regulatory functions rather than simple recycling functions.

Under pathological conditions, AEP has been shown to play key roles in the onset and progression of neurodegenerative diseases [9–14] and cancer [15–20]. In this context, the identification of its biological targets in neurodegenerative diseases, such as Parkinsońs and Alzheimeŕs, has allowed to understand its role in the onset and progression of these diseases at the molecular level, thus revealing AEP as a potential therapeutic target for the treatment of these conditions [9–14]. However, even though the connection between AEP overexpression in a plethora of human solid tumors and poor prognosis, increased malignancy, and worse overall survival has been reported [21–27], the mechanistic insights allowing to rationalize its role in the onset and progression of this disease are still lacking, hampering our ability to design novel, improved strategies for cancer treatment.

Here, we demonstrate that AEP plays a dual role in the resistance of breast cancer (BC) patients to genotoxic stress through the regulation of ATR and PPP1R10 levels, key DNA repair effectors [28, 29]. Specifically, AEP by suppressing ATR levels in BC cells increases genotoxic stress tolerance and reduces DNA damage-induced apoptosis, while maintaining proper cell cycle checkpoints allowing for efficient DNA repair by reducing PPP1R10 and protein phosphatase activity. Conversely, AEP deficiency results in BC cell death characterized by increased genomic instability and DNA damage signaling. Remarkably, in silico analyses using available clinical data from BC patients and further validated using an independent array of samples from IDBC patients showed that patients with AEP^high^/ATR^low^ levels can be efficiently identified as nonresponder patients, thus reinforcing the clinical significance of our findings and revealing AEP as a key contributor to tumor resistance. Besides, we also show that AEP inhibition sensitizes BC cells to cisplatin and etoposide treatment, further supporting a novel role for AEP in genotoxic stress tolerance in cancer cells and revealing AEP as a potential novel therapeutic target for the treatment of radioresistant BC cancer patients. Finally, our findings delineate a pan-cancer rule revealing a similar relationship between AEP^high^/ATR^low^ levels and reduced overall survival in other types of cancer. In summary, our data reveal AEP as a key factor contributing to cancer resistance, providing the rationale for the design of new strategies to treat resistant tumors by combining AEP inhibitors with current chemo- and radiotherapy approaches to sensitize them to genotoxic stress.

## Results

### AEP activity is required for cell cycle progression and cell division in cancer cells

AEP is overexpressed and correlates with poor prognosis and reduced overall survival in many human solid tumors. To explore this further we utilized the GEPIA2 online tool [30] and the TCGA database. These *in silico* analyses revealed that AEP mRNA expression levels were significantly higher (p<0.01) in a wide variety of human solid tumors compared to normal samples (Figures 1A-E, upper panels). Furthermore, survival analyses revealed that high AEP expression was statistically associated with poor prognosis and reduced overall survival in patients with these tumors (Figures 1A-E, bottom panels). Remarkably, in triple negative BC, high AEP expression levels correlated with a great reduction in overall survival, whereas patients with low AEP expression levels had almost one hundred percent survival. For this reason, we decided to focus on understanding the role of AEP in BC.

**Figure 1.**
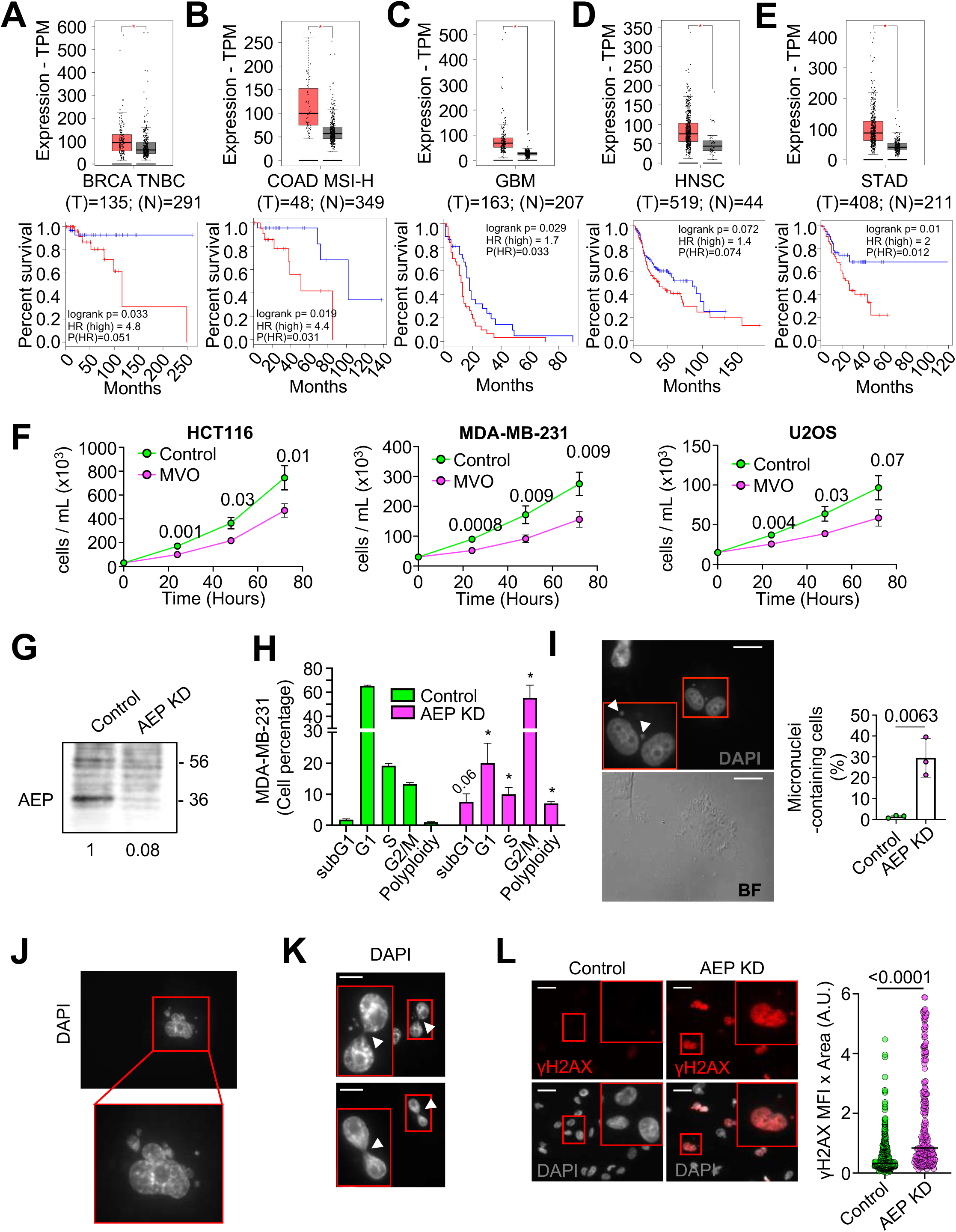
AEP deficiency reduces cell proliferation and impact cell cycle in cancer cells. Differential expression levels of AEP in different types of tumors were obtained using the GEPIA2 online application and extracted from The Cancer Genome Atlas (TCGA) database. Kaplan-Meier curves were constructed using the GEPIA2 online application to assess the correlation between AEP expression levels (low AEP blue line; high AEP red line) in different types of human tumors and overall survival. **(A)** AEP expression levels in BRCA triple-negative breast cancer (TNBC) patients (n=135, red) as compared to normal samples (n=291, grey) [upper panel] and Kaplan-Meier curves showing overall survival of patients with high (red line) or low (blue line) AEP expression levels [lower panel]. **(B)** AEP expression levels in colon adenocarcinoma with high genomic instability (COAD MSI-H) patients (n=48, red) as compared to normal samples (n=349, grey) [upper panel] and Kaplan-Meier curves showing overall survival of patients with high (red line) or low (blue line) AEP expression levels [lower panel]. **(C)** AEP expression levels in glioblastoma (GBM) patients (n=163, red) as compared to normal samples (n=207, grey) [upper panel] and Kaplan-Meier curves showing overall survival of patients with high (red line) or low (blue line) AEP expression levels [lower panel]. **(D)** AEP expression levels in head and neck squamous cell carcinoma (HNSC) patients (n=519, red) as compared to normal samples (n=44, grey) [upper panel] and Kaplan-Meier curves showing overall survival of patients with high (red line) or low (blue line) AEP expression levels [lower panel]. **(E)** AEP expression levels in stomach adenocarcinoma (STAD) patients (n=408, red) as compared to normal samples (n=211, grey) [upper panel] and Kaplan-Meier curves showing overall survival of patients with high (red line) or low (blue line) AEP expression levels [lower panel]. **(F)** Proliferation curves of different cancer cell lines (HCT116 [left panel], MDA-MB-231 [middle panel] and U2OS [right panel]) upon MVO-mediated AEP inhibition. Data represents average of 5 independent, biological replicas ± SD. **(G)** Immunoblot showing the shRNA-mediated AEP knock-down in MDA-MB-231 cells. **(H)** Cell cycle analyses of control and AEP shRNA-transduced MDA-MB-231 cells. Data represents average of 4 independent, biological replicas ± SD. * *p* value <0. 01. **(I)** Micrographs showing the presence of micronuclei in MDA-MB-231 cells upon shRNA-mediated AEP KD alongside quantitation. Data represents the average of 3 independent experiments, each one including more than 100 cell ± SD. Size bar = 27μm. **(J)** Micrographs showing examples of polyploid cells in shRNA-mediated AEP KD MDA-MB-231 cells. **(K)** Micrographs showing internuclear DNA bridges in shRNA-mediated AEP KD MDA-MB-231 cells. Arrowheads indicate DNA bridges. Size bar = 27μm. **(L)** Micrographs showing anti-γH2AX staining in both control (left panels) and AEP shRNA-transduced (right panels) MDA-MB-231 cells alongside quantitation representing mean fluorescence intensity (MFI). (n>200 cells). Size bar = 27μm.

Next, to characterize the role of AEP in cancer cells, we tested whether AEP inhibition with a highly specific inhibitor (MVO26630, hereafter referred to as MVO) [31] might affect cell proliferation. AEP inhibition significantly reduced cell proliferation in several cancer cell lines (Figure 1F). As previously reported [32], AEP inhibition resulted in a compensatory increase in AEP levels in cancer cells aimed at the recovery of its activity (Supplementary Figure 1A). Consequently, to inhibit this compensatory response, shRNA-mediated AEP knockdown (KD) approaches were used in MDA-MB-231 cells (Figure 1G). AEP KD resulted in increased cell death and an expansion of the G2/M phase (Figure 1H and Supplementary Figure 1D). Notably, a significant increase in both the percentage of micronuclei-containing cells (Figure 1I) and an increase in the number of polyploid cells (Figures 1H and 1J and Supplementary Figure 1D) in AEP KD MDA-MB-231 cells were observed, suggesting a potential role for AEP in suppressing genomic instability in BC cells. Additionally, AEP deficiency in cancer cells led to the formation of internuclear DNA bridges (Figure 1K) which were positive for γH2AX (Supplementary Figure 1I), consistent with an accumulation of DNA damage that led to chromosome segregation problems. In agreement with these observations, AEP deficiency resulted in an increase in γH2AX-positive cells even in the absence of external genotoxic insults (Figure 1L). Similar results were obtained for U2OS cells upon AEP KD (Supplementary Figure 1), further validating our observations in MDA-MB-231 cells. Taken together, these data suggest for the first time an unexpected role for AEP in chromosomal stability and DNA damage signaling/repair in cancer cells. However, the precise molecular mechanism (*i.e.*, the specific biological targets) by which AEP contributes to genomic stability in cancer remains unclear. Therefore, we undertook a high-throughput proteomic study to identify the biological targets and processes regulated by AEP, in order to gain insight into the mechanistic role of AEP in cancer cells.

### Chemical inhibition reveals noncanonical functions of AEP

As mentioned above, AEP inhibition frequently triggers a compensatory response [32, 33], making it difficult to identify potential AEP substrates. However, following a screening of different cell lines, we identified HEK293 cells as a cell line with high levels of AEP activity (Supplementary Figure 2A). Next, we verified that 16-hours MVO-treatment [31] effectively inhibited AEP activity in HEK293 cells without resulting in increased AEP activity (Supplementary Figure 2B). These data revealed HEK293 cells as appropriate candidates for further experiments designed to identify novel AEP targets that could allow to understand its role in chromosomal stability/DNA damage response in cancer cells.

Accordingly, to further our knowledge of the relationship between AEP and genomic stability we used label-free, quantitative high-resolution mass spectrometry on untreated and 24-hour MVO-treated HEK293 cells (Supplementary Figure 2C). The combined analyses of our proteomic data yielded 3,994 identified proteins, of which 283 statistically accumulated upon AEP inhibition (Figure 2A and Supplementary Table S1). The total number of peptides identified in the biological replicas was similar, with all replicates showing a Pearson correlation coefficient (r) value greater than 0.9 (Supplementary Figures 2D and 2E). Finally, gene ontology (GO) analyses of the 283 proteins significantly accumulating upon AEP inhibition (Figures 2A and 2C and Supplementary Table S1) revealed that 45.4% of these potential AEP targets localized to the nuclear compartment (Figure 2B), thus suggesting a role for AEP in the regulation of nuclear targets.

**Figure 2.**
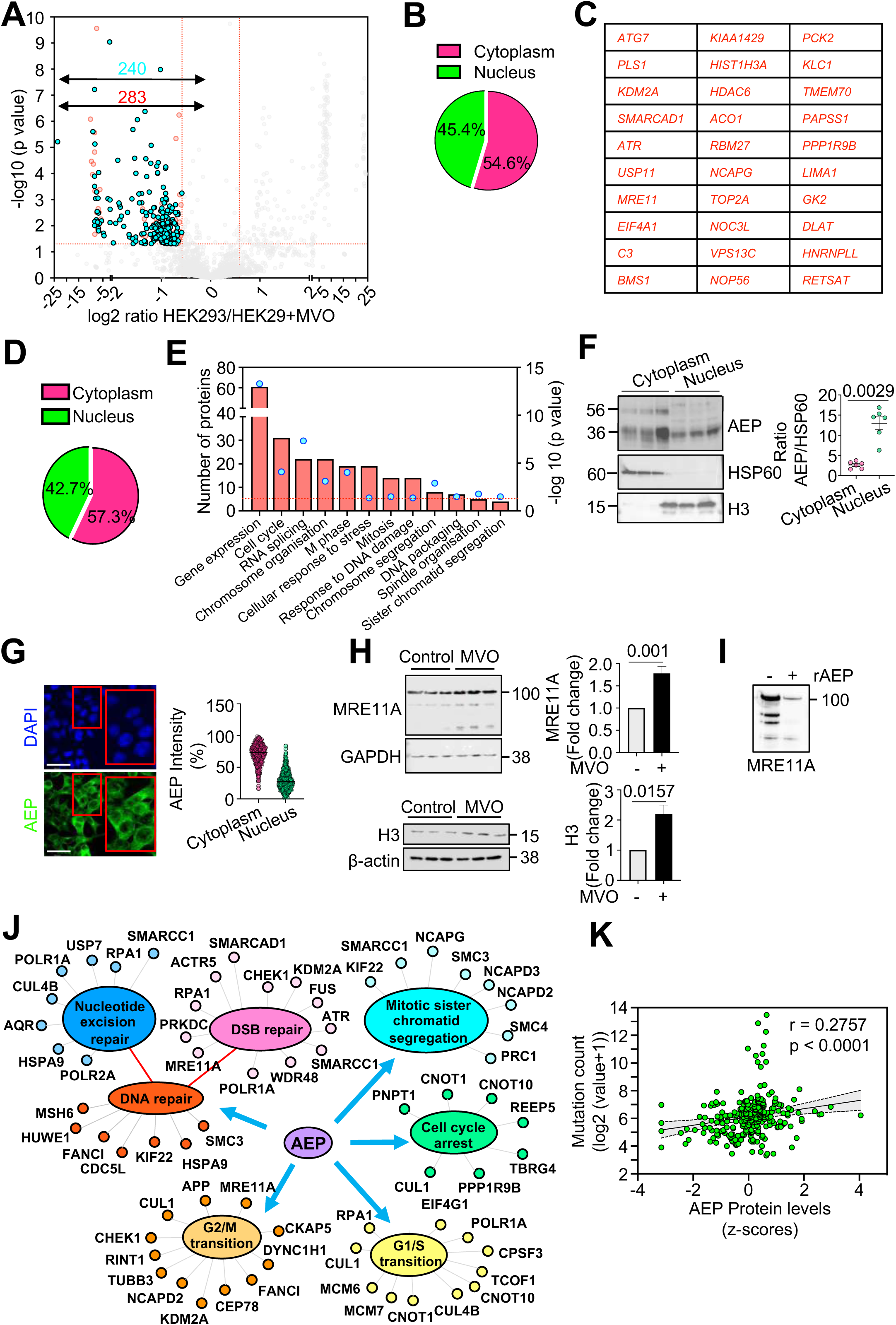
Proteomics study of MVO26630-mediated AEP inhibition in HEK293 cells reveals a putative nuclear role for AEP. **(A)** Volcano plot showing in red proteins whose levels are increased (283) and in cyan those that are increased and their putative cleavage site has been identified (240) upon 24-hours MVO26630 treatment. **(B)** Venn diagram showing nuclear and cytoplasmic distribution of the proteins accumulating upon MVO treatment. **(C)** Proteins showing the highest accumulation for which their putative AEP cleavage site has been identified upon MVO treatment as indicated by our proteomics data. **(D)** Venn diagram showing nuclear and cytoplasmic distribution of the proteins accumulating upon MVO treatment for which their putative AEP cleavage site has been identified. **(E)** GO analysis (Biological processes) of the proteins accumulating upon MVO treatment for which their putative AEP cleavage site has been identified (240 proteins). **(F)** Subcellular fractionation analysis of AEP localization in HEK293 cells alongside quantitation showing the average AEP/HSP60 signal ratio ± SD in 6 independent, biological replicas. **(G)** Micrographs showing AEP nuclear localization in HEK293 cells under normal conditions alongside quantitation of the AEP intensity normalized by the area per cell (n=635 cells), showing both cytoplasmic vs nuclear data (Size bar = 43.2um). **(H)** Immunoblot showing the levels of MRE11A and H3, AEP targets identified in our proteomics approach, in MVO-treated vs untreated HEK293 cells, alongside quantitation. Data represents the average of 3 independent, biological replicas ± SD. **(I)** Immunoblot showing the *in vitro* digestion of MRE11A overexpressed in HEK293 cells using recombinant AEP (rAEP) in pH6.5 for 3 hours. **(J)** Schematic representation of some of the newly identified targets of AEP highlighting the biological processes regulated by AEP as extracted from our GO analysis. **(K)** Correlation analyses between AEP protein expression levels (z-score) and mutation count (log2 (value+1)) in TCGA pan cancer samples obtained through cBioportal.

### Identification of novel AEP cleavage sites

As previously described, AEP is the only known protease that specifically hydrolyzes asparaginyl and, to a lesser extent, aspartyl peptide bonds [8]. Thus, an additional analysis of our proteomic dataset was conducted, focusing on tryptic peptides that are statistically accumulated upon AEP inhibition. These peptides were required to contain asparagine (Asn) and/or aspartate (Asp) residues in their sequence, as they could represent specific AEP cleavage sites. These analyses yielded 1,323 Asn/Asp-containing peptides statistically overrepresented upon AEP inhibition (Supplementary Table S2), corresponding to 994 proteins. This approach enabled the identification of potential AEP cleavage sites for 240 out of the 283 proteins that accumulated upon AEP inhibition (Figure 2A and Supplementary Tables S1-S2), thereby further reinforcing the potential role of AEP in the regulation of the levels of these proteins. Remarkably, some of the peptides identified through our proteomic analyses included already well-described AEP cleavage sites (Table 1 and Supplementary Table S2) [11, 34, 35], thus validating our experimental approach.

**Table 1.**
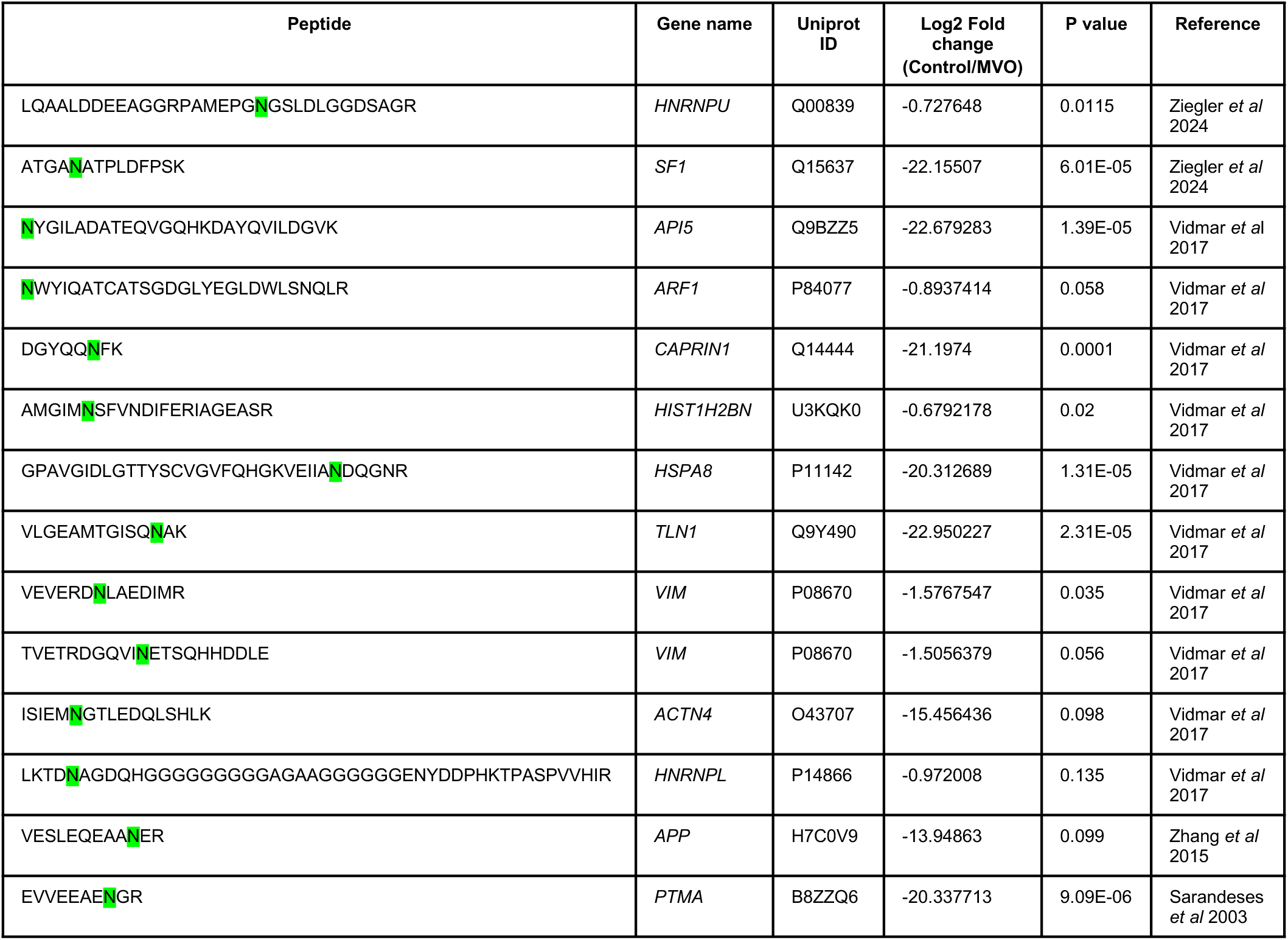
Peptides identified in our proteomics approach containing previously identified AEP cleavage sites (highlighted in green) indicating the accumulation in MVO treated samples as log2 fold change and the p value.

Interestingly, more than 40% of the proteins that accumulated upon AEP inhibition and that contained putative AEP cleavage sites, were localized to the nuclear compartment (Figure 2D). Moreover, a significant number of these proteins were associated with the DNA damage response, cell cycle regulation and mitosis/chromosome segregation (Figure 2E). Therefore, our combined proteomic data further support the hypothesis that AEP has a physiological function in the regulation of key nuclear proteins and, importantly, provides potential targets of AEP to interrogate at the molecular level its role in the regulation of these biological processes in cancer (Figure 1 and Supplementary Figure 1). Next, we decided to examine the subcellular localization and activity of AEP in HEK293 cells. Subcellular fractionation assays revealed that AEP was present in the nuclear fraction (Figure 2F). Furthermore, confocal microscopy analyses (Figure 2G) revealed that, while 70.6% ± 0.63 of AEP exhibited a clear cytoplasmic distribution, 29.4% ± 0.63 of AEP localized to the nuclear compartment. Finally, AEP activity in nuclear extracts that could be inhibited with MVO was detected using specific, fluorogenic substrates [32] *in vitro* (Supplementary Figure 2F). These results confirm the nuclear localization and activity of AEP and raise the possibility that a nuclear pool of AEP directly regulates the levels of these proteins.

Finally, we sought to validate our proteomic data by determining whether AEP inhibition resulted in the accumulation of targets identified through our proteomic approach. Some of the potential AEP targets were related proteins containing highly conserved potential cleavage sites (Supplementary Figure 2G). *In vitro* digestion of GFP-tagged versions of some of these proteins with recombinant AEP demonstrated its ability to completely digest these proteins (Supplementary Figure 2H), confirming that they are AEP substrates. Furthermore, AEP inhibition led to the accumulation of MRE11A and H3 (Figure 2H), both of which are among the highest ranked proteins identified as potential AEP targets in our proteomic approach (Figure 2C). Finally, *in vitro* assays revealed that recombinant AEP can digest overexpressed MRE11A (Figure 2I), thus reinforcing the idea that these proteins are bona-fide AEP targets. Thus, our combined data validate a potential role for AEP in regulating DNA damage signaling and genomic stability through the fine-tuning of the levels of these key proteins (Figure 2J). Notably, a pan-cancer data analyses from the TCGA database (n=263, including samples from breast, colorectal and ovarian epithelial cancers) revealed a positive correlation between tumor mutation count and the protein levels of AEP, further supporting a potential link between AEP and genomic stability (Figure 2K) and revealing the clinical significance of our findings.

### AEP activity controls ATR levels and negatively correlates with ATR in breast cancer patient survival

So far, our data obtained via both AEP chemical inhibition and genetic ablation suggest that AEP plays a role in cell cycle regulation, genomic stability, and DNA damage signaling in cancer cells (Figure 1). Furthermore, our proteomic approach strongly supports these observations by elucidating a previously unidentified function of AEP as a regulator of the levels of key nuclear proteins involved in the regulation of these biological processes in HEK293 (Figure 2). Therefore, to interrogate this new role of AEP in cancer cells, we analyzed its subcellular localization in BC cells. Both, confocal microscopy of endogenous AEP (Figure 3A) and subcellular fractionation analyses (Figure 3B) confirmed the nuclear localization of AEP in MDA-MB-231 cells. Remarkably, AEP is restricted to the cytoplasmic compartment in primary cells [32, 33], thus suggesting a pathological function for AEP in the nuclear compartment of cancer cells. Remarkably, similar results were obtained for colon carcinoma, osteosarcoma and glioblastoma cancer cell lines, further supporting the nuclear localization of AEP in cancer cells (Supplementary Figure 3A and 3B).

**Figure 3.**
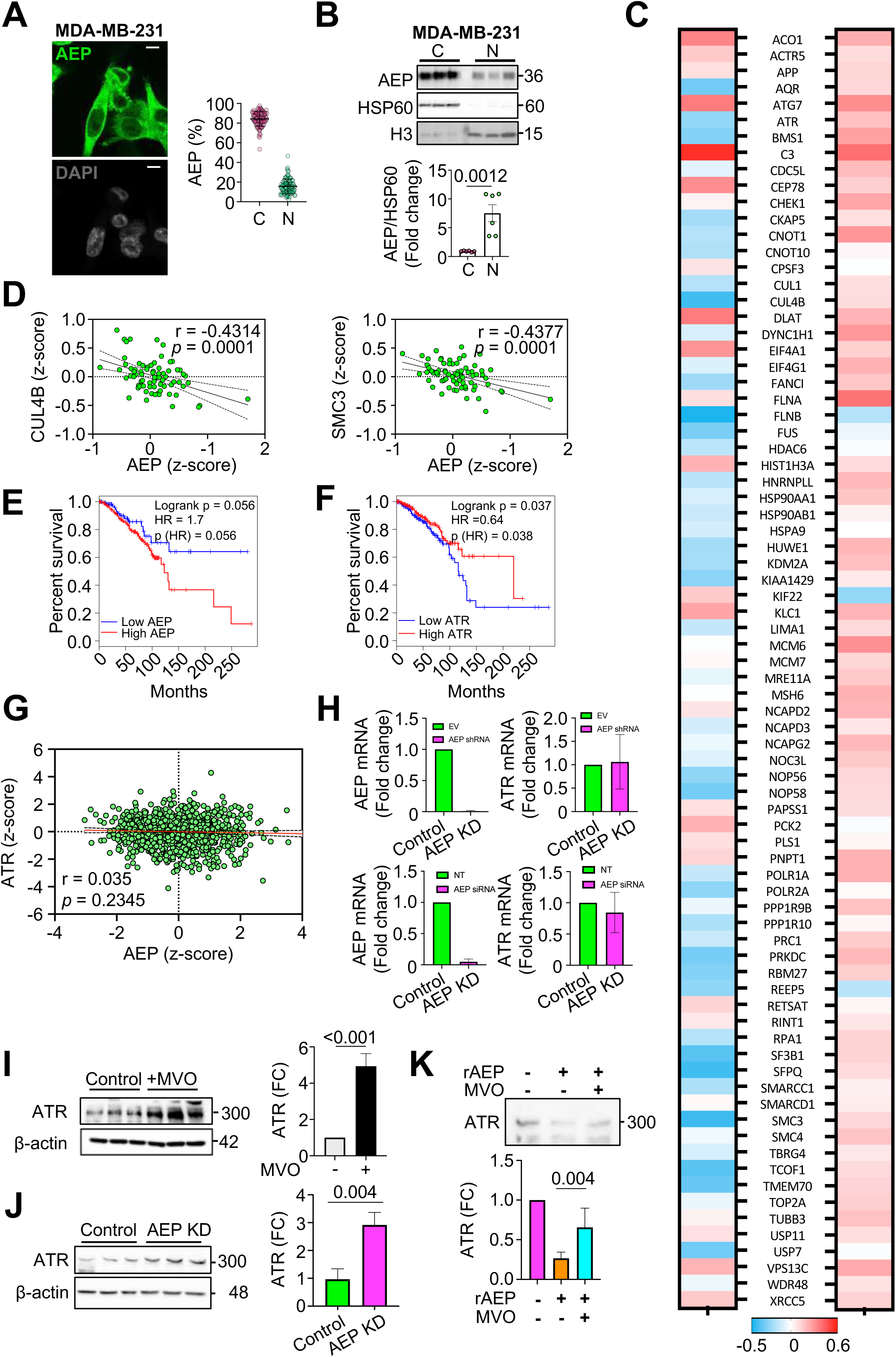
AEP localises in the nuclear compartment of cancer cells and regulates the levels of ATR. **(A)** Immunofluorescence showing the subcellular localisation of AEP in MDA-MB-231 (n>190 cells) alongside quantitation showing the cytoplasmic (C) and nuclear (N) intensity as percentage. (Size bar=6um). **(B)** Immunoblot showing the nuclear and membrane localisation of AEP in MDA-MB-231. Data represents the average ± SD of at six independent, biological replicas. **(C)** Correlation analyses between AEP and the main proteins identified as novel AEP targets in our proteomic experiment in breast cancer patients at the protein (left panel) and RNA (right panel) levels. **(D)** Correlation analysis of the protein expression levels of some of the novel AEP targets identified in our proteomic analyses (CUL4B and SMC3) vs AEP in BC patients using data obtained from the TCGA database. **(E)** Kaplan-Meier analysis of breast cancer patients expressing high (red line, n=503) or low (blue line, n=129) AEP levels. **(F)** Kaplan-Meier analysis of breast cancer patients expressing high (red line, n=319) or low (blue line, n=268) ATR levels. **(G)** Correlation analysis of the mRNA expression levels of ATR vs AEP in BC patients using data obtained from the TCGA database. **(H)** qPCR analyses of the mRNA expression levels of AEP and ATR in control (EV, empty vector) and AEP shRNA (upper panels) or control (NT, non-targeting) and AEP siRNA KD (bottom panels) MDA-MB-231 cells. **(I)** Immunoblot showing the effect of MVO-mediated AEP inhibition in the levels of ATR in MDA-MB-231 cells alongside quantitation. Data represents the average ± SD of at least three independent, biological replicas. **(J)** Immunoblot showing the effect of shRNA-mediated AEP knock-down in MDA-MB-231 cells in the levels of ATR alongside quantitation. Data represents the average ± SD of at least three independent, biological replicas. **(K)** Immunoblot showing the in vitro digestion of ATR using recombinantly expressed AEP alongside quantitation. Data represents the mean ± SD of six, independent biological replicas.

Next, we wanted to test whether the novel AEP targets identified in our proteomic approach were relevant in BC patients. To this end, correlation analyses using the TCGA BRCA dataset between the levels of expression of AEP and some of the proteins identified in our proteomic study as AEP targets and involved in DNA damage signaling and cell cycle (Figure 2, Supplementary Figure 2 and Supplementary Table S1) were carried out. Remarkably, increased AEP protein levels were found to be associated with a decrease in most of these proteins (Figure 3C, left panel), including key regulators of DNA damage signaling and cell cycle (Figure 3D and Supplementary Figure 3C), thus further supporting our proteomic data. Importantly, a similar correlation was not observed at the transcriptional level (Figure 3C, right panel), thus confirming a direct role for AEP in the regulation of the levels of these proteins not only in cell lines, but also in BC patients, further supporting our hypothesis of a novel, unexpected role of AEP in the regulation of genomic stability and DNA damage in BC.

As shown above, high AEP expression levels correlated with reduced overall patient survival in different types of cancer subtypes (Figures 1A-E), with AEP deficiency resulting in increased genomic instability, elevated γH2AX levels, and cell death (Figures 1H-L). Furthermore, as shown in the proteomic data (Figure 2 and Supplementary Table S1), AEP deficiency resulted in the accumulation of ATR, with reduced ATR protein levels observed in patients expressing high AEP protein levels (Figure 3C), further supporting a biologically relevant role for AEP in regulating ATR levels. Interestingly, ATR is one of the kinases involved in the phosphorylation of H2AX in response to DNA damage [36–38]. Therefore, we wondered whether AEP and ATR correlated with overall survival in BC patients including all the subtypes (TNBC, luminal A, luminal B and Her2+). Indeed, our analyses revealed that patients with high AEP expression levels had a poor prognosis and reduced overall survival (Figure 3E), whereas the opposite was observed for ATR, with low ATR levels correlating with reduced survival (Figure 3F). Remarkably, a similar observation was made in other types of cancer: high AEP expression levels (Supplementary Figure 3D, upper panels) and low ATR levels (Supplementary Figure 3D, bottom panels) correlated with poor prognosis and reduced overall survival, suggesting a more general role of the AEP/ATR axis in cancer.

Our analyses in BC patients showed a connection between AEP and ATR at the protein level (Figure 3C, left panel), that could not be observed at the transcriptional level (Figure 3C, right panel and Figure 3G). Moreover, qPCR analyses revealed that AEP KD in MDA-MB-231 cells, which was achieved via the use of either specific shRNAs (upper panels) or siRNAs (lower panels) against AEP, did not affect ATR expression levels (Figure 3H). These findings further strengthen the idea that AEP directly regulates ATR. Therefore, to further validate this connection we decided to study the effect of suppressing AEP in the levels of ATR in BC cells. Chemical inhibition and genetic ablation of AEP (Figure 3I and Figure 3J, respectively) resulted in a strong ATR accumulation in MDA-MB-231 cells, with similar results in U2OS cells (Supplementary Figure 3E), further validating a role for AEP in the regulation of ATR levels. In agreement with the finding that ATR is an AEP target, recombinant AEP was able to degrade purified ATR *in vitro* (Figure 3K). Overall, our data demonstrate AEP as a direct regulator of ATR protein levels in BC patients, thus identifying ATR as the molecular target for understanding the role of AEP as a regulator of DNA damage signaling and genomic stability in BC cells.

### AEP contributes to breast cancer resistance to genotoxic stress by reducing DNA damage signaling through the regulation of ATR and PPP1R10 levels

To advance our understanding of the AEP-ATR axis in BC at the molecular level, we explored the role of AEP deficiency in DNA damage signaling. First, we wondered whether the high levels of ATR observed in MDA-MB-231 cells upon AEP inhibition led to increased DNA damage signaling in response to genotoxic insults. Indeed, acute AEP inhibition resulted in increased DNA damage signaling in response to cisplatin treatment (Figure 4A), confirming the role of AEP in DNA damage signaling by reducing H2AX phosphorylation (γH2AX) through the regulation of ATR levels (Figures 2C, 3C, 3I-K).

**Figure 4.**
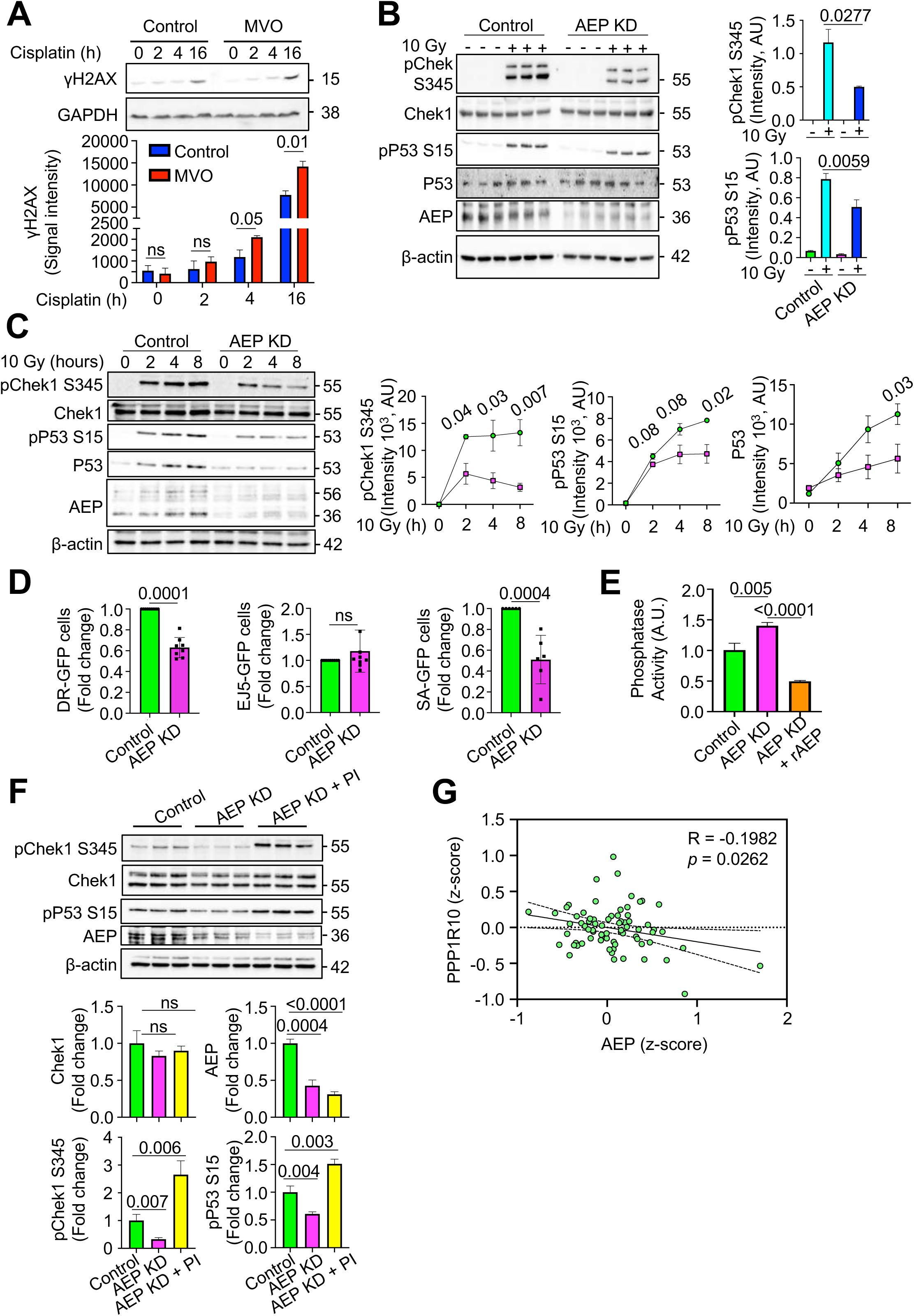
Nuclear AEP activity reduces DNA damage signalling while targeting phosphatase activity to maintain proper cell cycle checkpoint activation in breast cancer cells. **(A)** Immunoblot of γH2AX in MDA-MB-231 treated or not with MVO 50μM for 48 h and then stimulated with cisplatin 3μM, alongside quantitation. Data represents the mean + SD of three, independent biological replicas. **(B)** Immunoblot showing the phosphorylation levels of Chek1 and P53 upon 10 Gy radiation and 2 hours recovery in both control and AEP KD MDA-MB-231, alongside quantitation. Data represents the mean ± SD of three independent, biological replicas. **(C)** Time course showing the phosphorylation levels pf Chek1 and P53 upon 10 Gy radiation followed by 2-, 4- or 8-hours recovery, alongside quantitation in control (green circles) and AEP KD (pink squares) MDA-MB-231 cells. Data represents the mean ± SD of three independent, biological replicas. **(D)** Graph showing the effect of AEP deficiency in DNA repair using U2OS cells depleted or not of AEP and stably expressing DR-GFP, EJ5-GFP or SSA-GFP reporters constructs. After 72h GFP fluorescence was analysed by flow cytometry. Data show average fold change ± SD of the number of positive cells for each system compared to control cells. Data was generated from at least six independent, biological replicas. **(E)** Phosphatase activity in control vs AEP KD ± rAEP MDA-MB-231 cells. Data represents the mean ± SD of three independent, biological replicas. **(F)** Immunoblot showing the levels of phosphorylation of Chek1 upon 10 Gy radiation and 2 hours recovery in control MDA-MB-231 cells compared to AEP KD MDA-MB-231 pre-treated or not with phosphatase inhibitor (PI), alongside quantitation. Data represents the mean ± SD of three independent, biological replicas. **(G)** Correlation analysis of the protein expression levels of PPP1R10 vs AEP in BC patients using data obtained from the TCGA database.

In response to genotoxic insults, ATR phosphorylates Chek1 at serine 345, and subsequently pChek1 phosphorylates P53 at serine 15, thus inducing cell cycle arrest and allowing cells to repair DNA [39, 40]. Therefore, on the basis of our previous results and to further validate the connection between AEP and ATR in DNA damage signaling and chromosome stability, we hypothesized that the increase in ATR levels in AEP-deficient cancer cells (Figures 2C, 3C, 3I-K and Supplementary Figure 3E) could lead to increased pChek1 and pP53 levels, thus explaining the G2/M arrest observed in AEP-deficient cancer cells (Figure 1H). However, AEP deficiency in MDA-MB-231 cells resulted in reduced phosphorylation of both pChek1-S345 and pP53-S15 after 2 hours of recovery from a single dose of irradiation (10 Gy) (Figure 4B). To resolve this, we tested whether there were differences in the kinetics of phosphorylation and dephosphorylation of both pChek1 and pP53. Interestingly, upon irradiation, AEP-deficient MDA-MB-231 cells presented reduced levels of pChek1 and increased pChek1 dephosphorylation rates (Figure 4C, immunoblots and left panel), resulting in diminished levels of pP53 and total P53 accumulation (Figure 4C, immunoblots and middle and right panels).

Thus, AEP deficiency in cancer cells resulted in faster dephosphorylation rates of both Chek1 and P53 (Figure 4C) despite elevated ATR levels (Figures 3C and 3I-K). Remarkably, this unexpected result allowed to rationalize a previous observation. AEP-deficiency in cancer cells resulted in the accumulation of polyploid and micronuclei-containing cells, internuclear DNA bridges and elevated DNA damage resulting in cancer cell death (Figure 1 and Supplementary Figure 1). Specifically, the reduced activation of Chek1/P53 cell cycle checkpoint in response to irradiation observed in AEP KD cells (Figures 4B and 4C) explains why AEP deficiency in cancer cells resulted in genomic instability and cell death, due to their inability to properly activate Chek1/P53 cell cycle checkpoint in response to DNA damage.

Proper DNA damage repair requires the coordinated activation of ATR and the phosphorylation of H2AX to recruit the DNA repair machinery [38], but also the activation of cell cycle checkpoints to allow for DNA repair before the cell cycle can restart [41]. Furthermore, the inability to activate Chek1 in response to genotoxic stress and to induce cell cycle arrest to allow for DNA repair leads to aberrant mitosis, genomic instability, and cell death [42], features observed in AEP-deficient cancer cells (Figure 1). In this context, we hypothesized that AEP deficiency in cancer cells would result in reduced levels of DNA repair, thus explaining the genomic instability and cell death observed. To evaluate this, we measured the efficiency of the homologous recombination and nonhomologous end joining pathways using the DR-GFP [43], SA-GFP [44] and EJ5-GFP [44] reporter assays (Supplementary Figure 4). Indeed, shRNA-mediated AEP KD drastically reduced homology-directed repair, both classical homologous recombination (as measured with the DR-GFP system; Figure 4D, left panel) and single-strand annealing (as measured with the SA-GFP system; Figure 4D, right panel), without affecting nonhomologous end-joining (as measured with the EJ5-GFP system; Figure 4D, middle panel). Interestingly, homologous recombination contributes to genomic stability by facilitating DNA repair primarily during S and G2 phases [45], thus explaining the accumulation of DNA damage and increased genomic instability observed in AEP-deficient cancer cells. These data further support our previous observations and show that this new role of AEP is related to promoting genomic stability in BC cells (Figure 1) and patients (Figure 2K).

To further our understanding of the role of AEP in genotoxic stress tolerance in BC cells we aimed to investigate how AEP deficiency in BC cells resulted in reduced levels of both pChek1 and pP53, albeit increased levels of ATR. Our data, showing faster dephosphorylation of both pChek1 and pP53 in AEP-deficient cells, suggest that reduced phosphatase activity in BC cells could account for the prolonged activation of Chek1 and P53 shown in control MDA-MB-231 cells compared with AEP KD cells (Figure 4C), with previous works suggesting a potential role for AEP as a regulator of protein phosphatase activity [46–48]. In this context, AEP deficiency in MDA-MB-231 cells resulted in increased protein phosphatase activity, as measured with 6,8-difluoro-4-methylumbelliferyl phosphate (DiFMUP), which could be dampened by pretreating AEP KD samples with recombinant AEP (Figure 4E), thus supporting a role of AEP in cell cycle through the regulation of protein phosphatase activity in BC cells. Next, we wanted to evaluate whether protein phosphatase inhibition prior to irradiation in AEP-deficient BC cells could rescue Chek1 and P53 phosphorylation. Acute protein phosphatase inhibition in AEP-deficient MDA-MB-231 cells allowed for the recovery of both pChek1-S345 and pP53-S15 levels upon a single dose of irradiation (10 Gy) (Figure 4F), strengthen the idea that AEP functions as a regulator of pChek1 and pP53 through the regulation of protein phosphatase activity in cancer cells. Remarkably, our proteomic data showed that AEP inhibition results in a 2-fold accumulation of PPP1R10 (*p*=0.002) (Supplementary Table S1), a subunit of the PTW/PP1 complex involved in the regulation of the mitosis-to-interphase transition [49]. Interestingly, PPP1R10 genetic ablation results in prolonged Chek1 activation after ionizing-radiation-induced DNA damage [29], a phenotype that matches our observations in BC cells compared to AEP-deficient cells, thus revealing PPP1R10 as the potential AEP target explaining the increased protein phosphatase activity and the reduced levels of pChek1 and pP53 and increased genomic instability observed in AEP KD MDA-MB-231 cells. Furthermore, a negative correlation between AEP and PPP1R10 at the protein level is also observed in BC patients (Figures 3C and 4G), further reinforcing a direct role for AEP in the regulation of PPP1R10 levels, and therefore in the regulation of protein phosphatase activity in BC. These data further reinforce a role for AEP in regulating the activity of protein phosphatases, allowing BC cells to uncouple DNA damage signaling and cell cycle checkpoint activation, and provide the molecular basis that allow to understand the genotoxic tolerance-mediated by AEP in cancer cells.

In conclusion, our data suggest that, in BC cells, AEP contributes to genotoxic tolerance by reducing DNA damage signaling through a reduction in ATR levels, while promoting sustained pChek1 and pP53 phosphorylation to arrest the cell cycle and promote DNA repair through the regulation of PPP1R10 levels and protein phosphatase activity, thereby allowing cancer cells to scape DNA damage-induced cell death [50, 51].

### A high AEP-to-ATR protein ratio defines the response of patients with breast cancer to radiotherapy

Together, our data reveal a key role for AEP in BC resistance to genotoxic insults by proteolytically regulating ATR levels. Remarkably, despite the lack of correlation between AEP and ATR at the transcriptional level (Figures 3C, 3G and 3H), a negative correlation in the protein levels of AEP vs ATR could be observed in TCGA BC patients (Figures 3C and 5A), thus validating the findings in BC cells (Figures 3I and 3J). Interestingly, survival analyses using TCGA BRCA data demonstrated that patients with high AEP protein levels (AEP^high^, Figure 5B) or low ATR protein levels (ATR^low^, Figure 5C) had a worse prognosis and reduced overall survival, reinforcing their possible connection. However, to further investigate this novel AEP-ATR axis we analyzed whether AEP^high^/ATR^low^ patients presented differences in the overall survival compared with AEP^low^/ATR^high^ patients (Figure 5D). These analyses revealed that AEP^high^/ATR^low^ patients had reduced overall survival, whereas AEP^low^/ATR^high^ patients had one hundred percent overall survival (Figure 5E). Interestingly, AEP^low^/ATR^low^ or AEP^high^/ATR^high^ patients (Supplementary Figure 5A) did not significantly differ in overall survival (Supplementary Figure 5B), showing better overall survival than AEP^high^/ATR^low^ patients (Figure 5E), suggesting that low ATR or high AEP levels alone are not sufficient to explain the reduced overall survival.

**Figure 5.**
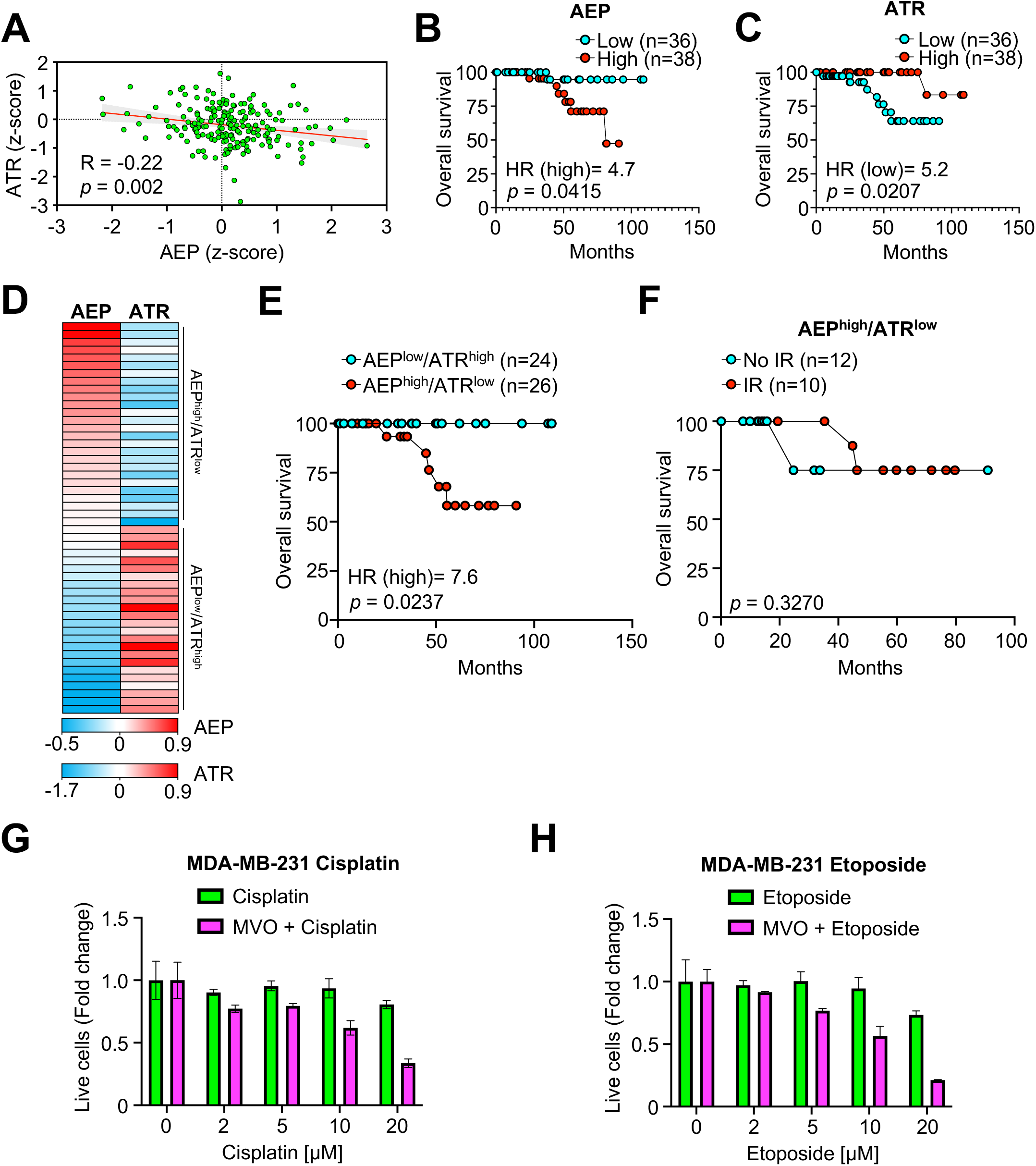
AEP contributes to breast cancer resistance to genotoxic stress. **(A)** Correlation analysis of AEP and ATR protein levels in BC patients using the TCGA breast carcinoma dataset. **(B)** Kaplan-Meier analysis in breast cancer patients expressing high (red dots) or low levels (cyan dots) of AEP at the protein level. **(C)** Kaplan-Meier analysis in BC patients expressing high (red dots) or low levels (cyan dots) of ATR at the protein level. **(D)** Heatmap showing the levels of protein (AEP and ATR, z-scores) expressed in BC patients. **(E)** Kaplan-Meier analysis in BC patients expressing AEP^low^/ATR^high^ levels (cyan dots) vs AEP^high^/ATR^low^ levels (red dots) at the protein level. **(F)** Kaplan-Meier analysis in BC patients showing AEP^high^/ATR^low^ protein levels treated (red dots) or untreated (cyan dots) with radiation. **(G)** Dose response of cisplatin in MDA-MB-231 cells in the presence (magenta bars) or absence (green bars) of MVO. **(H)** Dose response of etoposide in MDA-MB-231 cells in the presence (magenta bars) or absence (green bars) of MVO.

To explore the role of AEP in genotoxic tolerance in cancer, we investigated whether BC patients expressing different protein levels of AEP and ATR presented differences in the radiotherapy response. Patients with ATR^high^ levels had a good prognosis and no significant difference in overall survival, independent of whether they were treated or not with radiotherapy (Supplementary Figure 5C). Next, we focused on those patients expressing ATR^low^ levels, who showed reduced overall survival (Figure 5C). Remarkably, AEP^low^/ATR^low^ patients had a 100% percent overall survival rate when receiving radiotherapy, whereas untreated patients showed a reduced overall survival rate (Supplementary Figure 5E). This finding further corroborates that ATR^low^ levels in BC patients are not sufficient to explain tumor resistance to genotoxic stress. On the other hand, patients with AEP^high^ levels showed worse prognosis (Figure 5B), independent of whether they received radiotherapy or not (Supplementary Figure 5D), thus supporting a direct role for AEP in cancer resistance to genotoxic insults. Interestingly, while AEP^high^/ATR^high^ patients exhibited a favorable overall survival rate (Supplementary Figure 5B), the BC sample cohort with AEP^high^/ATR^low^ levels for whom data regarding radiotherapy treatment were available (n=22), accounted for most of the reduced overall survival observed in AEP^high^ expressing patients (Figure 5B), showing no significant difference independent of their being treated or not with radiotherapy (Figure 5F). These data confirm the role of AEP in the regulation of tumor resistance to genotoxic stress through the regulation of ATR levels, with AEP^high^/ATR^low^ patients showing a limited response to radiotherapy. On the contrary, all the remaining patient groups had good overall survival and response to radiotherapy. Hence, our data reinforce a key role for this novel AEP-ATR axis in genotoxic tolerance in BC, suggesting the possibility of combining AEP inhibitors with current radiotherapy or chemotherapy approaches to increase treatment efficiency in AEP^high^/ATR^low^ patients.

Finally, to test our hypothesis, we studied whether AEP inhibition in BC cells sensitizes them to genotoxic insults. This study revealed that 24-hour AEP inhibition in MDA-MB-231 cells sensitized them to the chemotherapeutic drugs cisplatin and etoposide, leading to increased cell death, further corroborating the role of AEP in BC cells resistance to genotoxic stress (Figures 5G and 5H). Remarkably, similar findings were obtained in U2OS cells (Supplementary Figures 5F and 5G), in which we previously showed that AEP-deficiency also resulted in cell death (Supplementary Figure 1) and increased ATR levels (Supplementary Figure 2), thus further validating our hypothesis and the novel role of AEP as a regulator of ATR levels and as therapeutic target to sensitize cancer cells to genotoxic insults.

### Increased nuclear AEP levels reduce ATR nuclear accumulation in nonresponder invasive ductal breast carcinoma patients

Our previous data (Figure 5 and Supplementary Figure 5) suggest that increased AEP levels are linked to reduced radiotherapy response. Therefore, we propose that AEP could be a determinant of the level of radiotherapy resistance in BC patients. To validate whether AEP could discriminate the response to therapy, we analyzed the levels of AEP in a series of human IDBC patient samples previously classified as responders (n=10) and nonresponder (n=10). Immunohistochemical analyses revealed that AEP was expressed mainly in ductal BC cells (Figure 6A and Supplementary Figure 6A), with immunofluorescence approaches revealing higher AEP levels in ductal BC cells from nonresponder patients compared to responder patients (Figure 6B). Next, we investigated whether a negative correlation between AEP and ATR levels could also be detected in these cancer cells. Indeed, higher AEP levels in nonresponder patients correlated with reduced ATR nuclear levels in 9 out of 10 patients (Figure 6C), further confirming our previous observations revealing a direct role for AEP in regulating the levels of ATR and reinforcing their connection with tumor resistance to radiotherapy. Conversely, responder patients included those with AEP^low^/ATR^high^ levels or similar AEP-to-ATR levels (Figure 6C), further reinforcing our previous observation that AEP^high^/ATR^low^ patients exhibited the poorest response to radiotherapy. Moreover, our analysis confirmed the AEP/ATR ratio as an efficient predictive biomarker for the stratification of responder (AEP^low^/ATR^high^, AEP^low^/ATR^low^ or AEP^high^/ATR^high^) and nonresponder (AEP^high^/ATR^low^) patients (Figure 6D), further supporting an unexpected connection between AEP and ATR levels in genotoxic tolerance in cancer cells (Figure 5). Moreover, our previous data indicate that AEP specifically accumulates in the nucleus of cancer cells; thus, we wondered whether AEP nuclear localization could also be observed in primary tumor samples supporting the idea that a nuclear pool of AEP specifically targets and fine-tunes ATR nuclear levels. Indeed, increased AEP nuclear levels were specifically detected in ductal BC cells from nonresponder patients (Figure 6E). Furthermore, an increased nuclear AEP/ATR ratio (Figure 6F and Supplementary Figure 6B) was observed in nonresponder patients. In this context, on average, 60% of nonresponder patients had ductal carcinoma cells with a nuclear AEP/ATR ratio greater than 0.5, whereas 16% of responder patients did. These combined data stresses the clinical relevance of the direct role of AEP in the fine-tuning of ATR levels in BC cells and in their resistance to radiotherapy.

**Figure 6.**
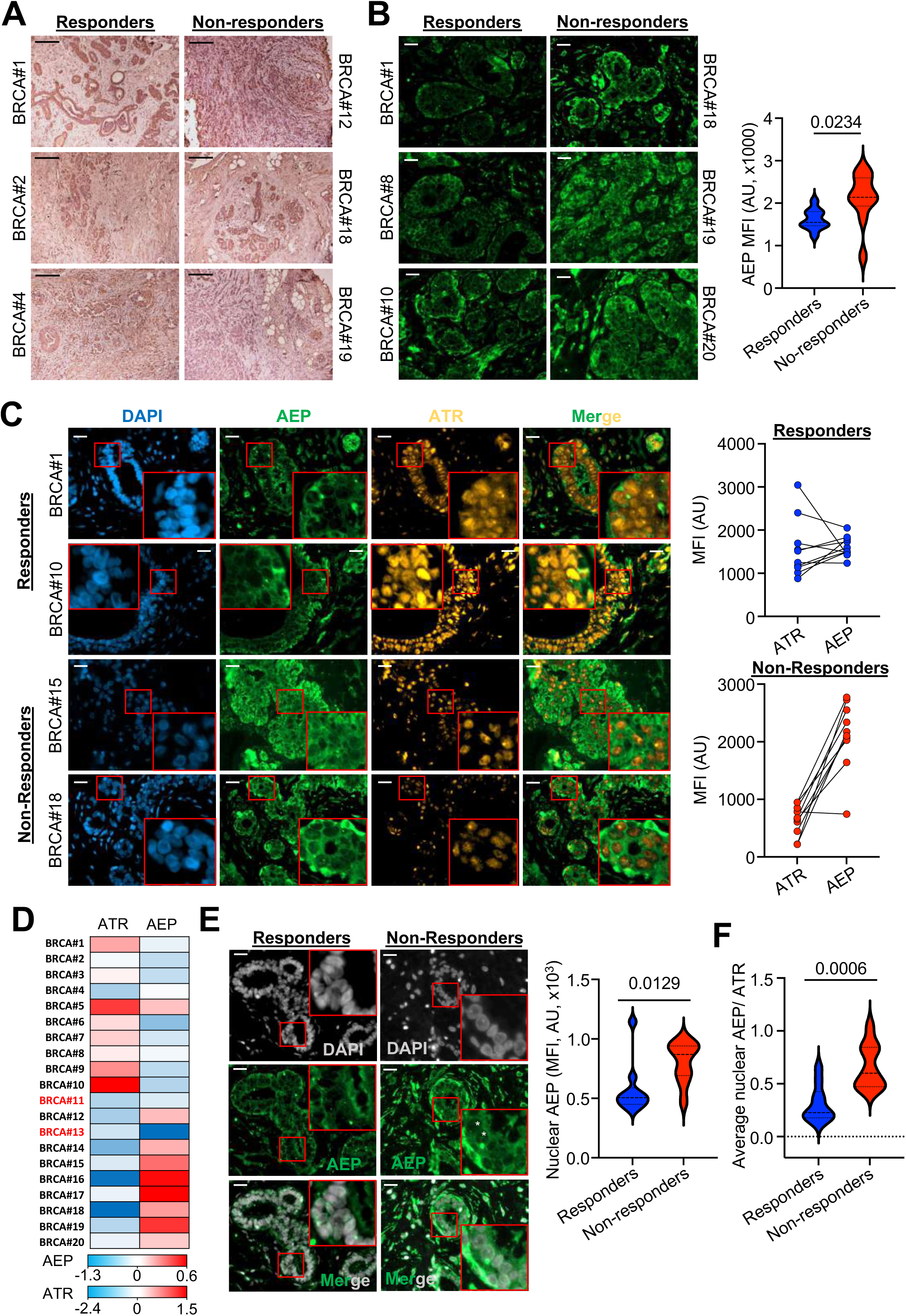
Increased AEP levels reduce ATR nuclear levels in ductal breast carcinoma cells in chemotherapy non-responder human patients. **(A)** Microscopy images (10x) of the immunohistochemical analysis of AEP expression in responder and non-responder invasive ductal breast carcinoma samples. (Size bar=200μm). **(B)** Immunofluorescence analysis of AEP expression levels in ductal breast carcinoma cells in responder and non-responder patients, alongside quantitation. Data represents the average AEP expression levels quantified using samples from 10 responder (50 cells per patient) and 10 non-responder (50 cells per patient) patients. (Size bar = 20μm). **(C)** Immunofluorescence analysis of the expression levels of AEP and ATR in ductal breast carcinoma cells in responder and non-responder patients. Each pair represents the average ATR and AEP expression levels obtained upon quantification of data obtained per patient using 50 cells per patient. (Size bar = 20μm). **(D)** Heatmap showing the protein levels (AEP and ATR, z-scores) expressed in invasive ductal breast cancer patients organized into responder and non-responder patients. **(E)** Immunofluorescence analysis of the nuclear AEP expression levels in ductal breast carcinoma cells in responder (n=10) and non-responder (n=10) patients, alongside quantitation. Data represents the average nuclear level of AEP obtained from the quantification of at least 80 cells per patient. (Size bar = 20μm). White asterisk indicates AEP-positive nuclei in non-responder samples as compared to responder samples. **(F)** Nuclear AEP/ATR ratio obtained from responder (n=10) and non-responder (n=10) patients. Each dot represents the average nuclear AEP/ATR ratio obtained for each patient upon quantification of at least 80 cells per patient.

## Discussion

Understanding tumor resistance and identifying novel druggable targets to sensitize them to radio- or chemotherapy approaches is a major research focus in oncology. This is because, despite current advancements in both strategies that have contributed to improve treatment outcomes [1], cancer resistance still accounts for 80-90% of deaths in cancer patients [2, 3].

AEP overexpression correlates with poor prognosis, increased malignancy, and worse overall survival in many human solid tumors [16, 19, 20, 22, 23, 25, 52, 53]. Remarkably, novel roles of AEP in tumor onset and progression are emerging, with the identification of the molecular targets of AEP allowing to understand its role in cancer biology. Recent work demonstrates that AEP suppresses lung metastasis, suggesting a potential role for AEP in BC onset and progression [17]. Furthermore, AEP promotes glioblastoma tumorigenesis and proliferation through p53 inactivation [16], facilitates tumor malignancy through TMOD3 processing [54], and mediates cancer cell adaptation to harsh environments through DDX3X cleavage [18]. Together, these studies strongly support a key role for AEP in cancer, highlighting the importance of identifying novel targets and biological processes regulated by AEP contributing to the onset and progression of this disease. Here, we reveal AEP as a determinant of radiotherapy resistance in BC. Our data show that AEP accumulates in radiotherapy nonresponder invasive breast carcinoma patients, playing a dual role in tumor resistance to genotoxic stress. Moreover, we reveal AEP as a previously unidentified predictive biomarker for radioresistant BC patients and as a therapeutic target for sensitizing BC cells to current radiotherapy and chemotherapy approaches.

At the mechanistic level, we demonstrate that AEP functions as a determinant of genotoxic stress tolerance in BC by uncoupling DNA damage signaling from cell cycle checkpoints and DNA damage repair. First, our work shows that AEP suppresses ATR levels, thus reducing DNA damage-induced apoptosis in response to genotoxic stress [55–58]. Importantly, reduced ATR levels in BC patients have been shown to correlate with reduced overall survival [59]. Furthermore, ATR haploinsufficiency resulting in reduced ATR activity has been revealed as a driver of tumorigenesis [60–64]. In this context, reduced ATR levels are tumor-prone due to partial DNA damage response defects leading to increased genomic instability [62]. However, the mechanisms controlling ATR levels remain unclear. Our work directly elucidates AEP as a previously unidentified regulator of ATR levels in cancer, thus allowing cancer cells to better cope with genotoxic stress. On the other hand, we show that AEP reduces PPP1R10 levels, leading to sustained Chek1 and P53 phosphorylation, thus promoting the activation of the G2/M checkpoint and facilitating DNA damage repair in response to genotoxic insults in BC. Interestingly, genetic ablation of PPP1R10, a regulatory subunit of PP1 [49], results in prolonged Chek1 and P53 activation after ionizing-radiation-induced DNA damage [29], a phenotype that matches our observations in BC cells compared to AEP-deficient cells. Furthermore, reduced PP1 levels have been shown to promote tumorigenesis and to correlate with reduced overall survival [65–68], however the underlying mechanisms controlling PP1 activity in BC cells remain unclear. Our study reveals AEP as a regulator of cell cycle progression in response to genotoxic stress through the modulation of PPP1R10 levels, thus explaining the reduced levels of pChek1 and pP53, reduced DNA damage repair, and increased genomic instability observed in AEP KD MDA-MB-231 cells. Taken together, our findings provide the underlying mechanisms to understand the role that AEP plays in tumor resistance against genotoxic stress.

The clinical significance of our findings is demonstrated by our analyses revealing that radioresistant BC patients are efficiently identified as AEP^high^/ATR^low^ BC patients. First, we reveal that AEP accumulates in ductal breast carcinoma cells from nonresponder patients, resulting in reduced ATR nuclear levels, with these patients showing a higher AEP/ATR ratio. Conversely, AEP^low^/ATR^high^, AEP^low^/ATR^low^ or AEP^high^/ATR^high^ patients are efficiently discriminated as responders. Thus, our work identifies for the first time the AEP/ATR axis as a predictive biomarker for the identification of radiotherapy nonresponder patients. Moreover, we demonstrate that AEP inhibition in BC cells sensitizes them to chemotherapeutic reagents such as cisplatin or etoposide, revealing AEP as a promising therapeutic target for the treatment of radioresistant BC patients. Importantly, our data reveal a similar relationship between AEP and ATR levels not only in BC, but also in other types of cancer. Specifically, our findings indicate that in cancers where high AEP expression levels are associated with poor prognosis, the opposite is observed for ATR. Thus, these observations suggest a more general role for this newly identified AEP/ATR axis in cancer, although the connection between the AEP/ATR axis and tumor resistance in other types of cancer remains to be explored. The present study reveals the AEP/ATR/PPP1R10 axis as a determinant of genotoxic stress tolerance and radiotherapy resistance in BC and provides the theoretical basis for the design of novel AEP-targeted therapies in the treatment of chemotherapy or radiotherapy resistant BC tumors.

## Conclusion

In the current study we identify the role of the AEP/ATR/PPP1R10 axis in genotoxic stress tolerance in breast cancer patients. We demonstrate that AEP suppresses ATR levels resulting in reduced levels of DNA damage signaling, thus allowing BC cells to scape DNA damage-induced cell death. Besides, we reveal AEP reduces protein phosphatase activity through the suppression of PPP1R10 levels in BC, thus allowing cell cycle checkpoint activation and efficient DNA damage repair. Therefore, our work uncovers a dual role for AEP in genotoxic stress resistance in BC by uncoupling DNA damage response from cell cycle checkpoint activation (Figure 7). Furthermore, we provide strong evidence supporting AEP as both, a novel therapeutic target and an efficient predictive biomarker for the treatment and stratification of nonresponder patients.

**Figure 7.**
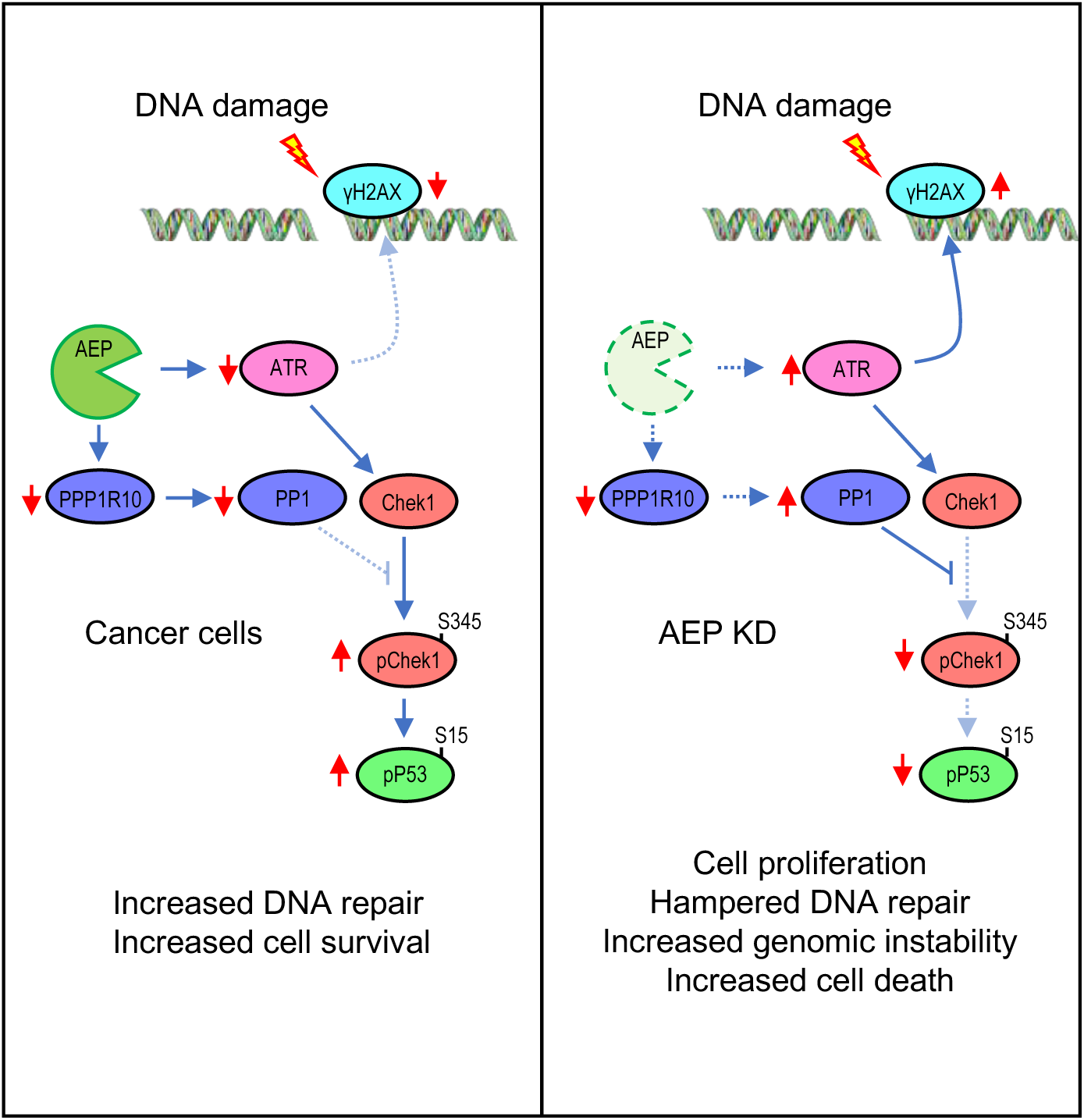
AEP plays a dual role in radiotherapy resistance in BC patients. In BC patients, AEP reduces DNA damage response by suppressing ATR levels, while maintaining sustained Chek1 and P53 activation through the suppression of PPP1R10 levels and PP1 activity, thus allowing cancer cells to scape DNA damage-induced cell death and to efficiently repair DNA damage, thus explaining the role of AEP in radiotherapy resistance (left panel). Conversely, AEP deficiency in cancer cells result in increased ATR levels leading to elevated DNA damage signaling and increased PP1 activity, resulting in reduced levels pf pChek1 and pP53 leading to reduced DNA repair, increased genomic instability and cell death (right panel).

## Materials and Methods

### Human tumor samples

A total of 20 paraffin-embedded human breast cancer (BC) samples at diagnosis were provided by the Biobank of the Anatomy Pathology Department (record number B.0000745, ISCIII National Biobank network) of the MD Anderson Cancer Centre Madrid, Spain. In summary, the patient samples were selected according to pathological criteria as infiltrating ductal breast carcinoma diagnosed as G2-3. Patients, aged 49+8 years without previous oncological diseases, received standard breast cancer treatment and radiotherapy as appropriate[69]. The response to radiotherapy was assessed according to established criteria[70]. This study was conducted in accordance with the ethical standards set forth in the Spanish legislation (Ley de Investigacion Organica Biomedica, 14 July 2007) and was approved by the ethical committees of the MD Anderson Cancer Centre Madrid, Spain. A comprehensive written informed consent was obtained from all participants.

### Cell culture

Cell lines were acquired from the American Type Culture Collection (ATCC). HEK293 cells (RRID:CVCL_0045) were cultivated in low-glucose DMEM (Sigma, #5546), supplemented with 100U/ml penicillin, 100μg/ml streptomycin and 2mM L-glutamine. Cancer cell lines (*i.e.,* HCT116 (RRID:CVCL_0291), MDA-MB-231 (RRID:CVCL_0062), U2OS (RRID:CVCL_0042), U87 and T98) were cultured in high-glucose DMEM containing 100U/ml penicillin, 100μg/ml streptomycin, 2mM L-glutamine. All cell lines were cultured at 37°C in an atmosphere of 5% CO2.

### Antibodies

All antibodies used in this manuscript were used a 1:1000 dilutions for immunoblotting or 1:200 dilutions for immunofluorescence. Antibodies against GAPDH (clone 14C10, #2118), COXIV (clone 3E11, #4850), β-actin (clone 13E5, #4970), P53 (clone 1C12, #2524), pS15-P53 (#9284) and pS345-Chek1 (clone 133D3, #2348) were purchased from Cell Signaling. HSP60 (clone EP1006Y, #ab45134) and SFPQ (#ab38148) were obtained from abCam. Antibodies against H3 (#SAB4200651) and pS139-H2AX (clone JBW301, #05-636) were acquired from Sigma. Antibodies against GFP (#A-11122), KU80 (clone 9403, #MA1-23314) and ATR (clone 2B5, MA1-23158) were obtained from Thermo Scientific. Antibody against MRE11A (#203501-T38) was acquired from SinoBiologicals. Antibodies against CHEK1 (clone G4, #sc-8408) and TOP2A (clone F12, #sc-365916) were obtained from SantaCruz. Horseradish peroxidase-conjugated streptavidin (SAV-HRP, #016-030-084) was obtained from Jackson Immunoresearch. Sheep anti-AEP was obtained as previously described[71] and sheep anti-CtsL and CtsH were generous gifts from Drs Tina Zavašnik-Bergant and Janko Kos (Jožef Stefan Institute, Ljubljana, Slovenia).

### Tissue immunohistochemistry and immunofluorescence in human samples

Paraffin-embedded samples were submitted to standard immunohistochemistry (IHC) protocol. Briefly, after deparaffinization and hydration, antigen retrieval was performed with 10 mM citrate buffer pH 6.0 in microwave, followed by peroxidase inhibition with 3% H2O2 and blocking of unspecific binding with 5% BSA. Incubation with primary antibody anti-AEP 1:100 at 4°C overnight followed by secondary anti-sheep-HRP for 1h at RT. DAB reaction kit (catalogue no. SK-4100) was used to visualize the peroxidase reaction product. Samples without the primary antibody were used as a negative control. All the sections were counterstained with hematoxylin. Images were obtained with a Leica microscope and Leica digital camera. For immunofluorescence staining the peroxidase inhibition step was omitted. After blocking, sections were incubated with anti-AEP (as mentioned above) and anti-ATR 1:200 at 4°C overnight, followed by anti-sheep and anti-mouse Alexa Fluor-conjugated secondary antibodies, respectively. Sections analyzed and images acquired with a Zeiss ApoTome 3 microscope.

### Enzymatic activity assays

Total cell lysates were prepared in 0.2M Na citrate buffer pH 4.0 containing 1% Triton X-100 (Sigma, #T9284). AEP activity was determined using 20μg of total protein in 200μl of assay buffer (0.2 M Na citrate buffer pH 4.0, supplemented with 1mM DTT) containing 10μM of AEP substrate (Z-Ala-Ala-Asn-7-amino-4-methylcoumarie (AMC), (Bachem, #4033201)). Release of AMC was quantified at 460nm in a fluorescence plate reader (Fluorostar Optima, BMG Labtech).

### In vitro digestion assay

Recombinantly expressed proteins were extracted in 50mM sodium acetate buffer containing 1% Triton X-100, containing 4mM EDTA, pH6.5 and supplemented with protease inhibitor cocktail (Roche). Equal amounts of protein (20μg) were incubated in a pH-modified AEP reaction buffer (50mM sodium acetate, 4mM EDTA, pH6.5, 5mM DTT and protease inhibitor cocktail) for 1 hours at 37°C in the presence or absence of recombinant AEP.

### Phosphatase activity assay

Total cell lysates from control and AEP deficient MDA-MB-231 cells were prepared in Tris-HCl 50mM, pH 8 buffer, containing 100 mM NaCl, 2mM EDTA and 1% NP-40, and supplemented with Protease Inhibitor Cocktail and PMSF. All samples were incubated at 37°C for 30 minutes in the presence or absence of recombinantly expressed AEP, prior to the measurement of phosphatase activity. Phosphatase activity was measured using DiFMUP (6,8’-Difluoro-4-Mrthylumbelliferyl Phosphate) following manufacturer instructions.

### Immunofluorescence microscopy

Cells (5 x 104 cells/coverslip) were fixed in PBS containing 4% paraformaldehyde for 20 min at room temperature. Then, cells were washed with PBS three times, permeabilized with 0.2% Triton X-100 in PBS for 5 minutes at room temperature, washed again three times with PBS and incubated in blocking solution (1% BSA in PBS) for 20 min at room temperature. Then, cells were stained using a 1:200 dilution of primary antibody in blocking solution for 45 min at room temperature, washed three times with PBS and stained with a 1:500 dilution of the Alexa-488 or Alexa-594 conjugated secondary antibody (Invitrogen). Finally, coverslips were washed three times in PBS and mounted using Prolong Gold Antifade with DAPI (Thermo Scientific) and images acquired using a LSM700 confocal microscope (Leica) or a Zeiss Apotome 3. Quantitative analyzed of the immunofluorescence images were done using ImageJ (RRID:SCR_003070) software.

### DNA constructs

The pGL4.47[luc2P/SIE/Hygro] vector (Promega, #E4041) was used to measure STAT3 upon IL6 stimulation of AEP inhibition using different specific inhibitors (*i.e.*, MVO26630, WL-864 or WL-1030) activity in HEK293 cells transformed using JetPEI (Polyplus, #101000053), following manufacturer instructions. DNA sequences coding for Nucleolar Protein 56 (NOP56, #HG16916-U), Nucleolar Protein 58 (NOP58, #HG20632-U), DNA replication licensing factor MCM6 (MCM6, #HG14317-G), DNA replication licensing factor MCM7 (MCM7, #HG16465-G), Heat shock protein HSP 90 alpha (HSP90AA1, #HG11445-G) and Heat shock protein HSP 90 beta (HSP90AB1, #HG11381-G) were obtained from SinoBiological, cloned into pEGFP-N1 (Clontech, #6085-1) and transformed into HEK293 (RRID:CVCL_0045) cells using JetPEI following manufacturer protocol. DNA coding for asparagine endopeptidase (AEP) was cloned into pSEMS-meGFP[72].

### Sample preparation for label-free mass spectrometry

HEK293 cells were treated or not with 50μM MVO26630 for 24 h. Next, cells were washed twice with ice-cold PBS, scrapped from cell culture dishes and collected by centrifugation at 300 g for 5 min at +4°C, and resuspended in SDS-containing lysis buffer (1% SDS in 100mM Triethylammonium Bicarbonate buffer (TEAB, Sigma, #T7408)), kept on ice for 10 min to guarantee cell lysis. Then, cell lysates were centrifuged at 20,000 g for 10 min at +4°C and the supernatant transferred to a clean low-protein binding tube. Protein concentration was determined using the BCA Protein Assay Kit (Thermo Scientific, #23227), and 1 mg of protein per experiment was reduced with 10mM DTT (Sigma, #D0632) for 1 h at 55°C and then alkylated with 20mM iodoacetamide (IAA, Sigma, #6125) for 30 min at RT. Protein was then precipitated overnight at -20°C using six volumes of chilled (-20°C) acetone. After precipitation, protein pellet was resuspended in 1ml of 100mM TEAB and digested with Trypsin (1/100 w/w, Thermo, #90058) and digested overnight. Finally, digested samples were cleared by centrifugation at 20,000g for 30 min at +4°C, and peptide concentration quantified with a Quantitative Colorimetric Peptide Assay Kit (Thermo, #23275).

### Mass spectrometry

Peptide samples were analyzed using a nanoflow liquid chromatography system (Ultimate 3000 RSLCnano system, Thermo Scientific) coupled to an LTQ Orbitrap Velos Pro mass spectrometer (Thermo Scientific). Samples, usually 10μl, were loaded onto a C18 trap column and washed for 5 min with 0.1% formic acid. The peptides were resolved using a gradient (130 min, 0.3μl/min) of buffer A (0.1% formic acid) and buffer B (80% acetonitrile in 0.08% formic acid): 2% buffer B for 4 min, 2%-40% buffer B for 64 min, 40%-98% buffer B for 2 min, 98% buffer B for 15 min, 98%-2% buffer B for 1 min and 2% buffer B for 44 min. Peptides, initially trapped on an Acclaim PepMap 100 C18 column (100μm x 2cm, Thermo Scientific), were then separated on an Easy-Spray PepMap RSLC C18 column (75μm x 50cm, Thermo Scientific), and finally transferred to a LTQ Orbitrap Velos Pro mass spectrometer via an Easy-Spray source set at 50°C and a source voltage of 1.9 kV. For the identification of the peptides, a top 15 method (1 MS plus 15 MS^2^, 100 min acquisition) consisting of full scans and mass range (m/z) between 335 and 1,800 (m/z). The Orbitrap was operated in a profile mode, resolution of 60,000 with a lock mass set at 445.120024 and a max fill time of 500ms. LTQ was operated in a centroid mode with isolation width = 2.00 (m/z), normalized collision energy = 35.0, activation time = 10ms and max fill time of 100ms.

### Mass spectrometry analyses

LTQ Orbitrap Velos Pro.RAW files were analyzed, and peptides and proteins quantified using MaxQuant[73]. All settings were set as default, except for the minimal peptide length of 5, and Andromeda search engine was configured for the Uniprot Homo sapiens protein database. Peptide and protein ratios only quantified in at least two out of the three independent, biological replicas were considered. Statistical significance was calculated using Student’s t test.

### Gene Ontology Analyses

Gene Ontology (GO) analyses to reveal statistically over-represented terms were performed using ClueGo[74]. The proteins accumulating upon AEP inhibition or biotinylated when using AEP-TiD in our mass spectrometry analyses were used as a test dataset and a reference set of GO annotations for the human proteome was used to carry out the GO enrichment analyses. Analyses were done using a right-sided hypergeometric test and only GO terms (*i.e.,* cellular location and biological process) with p < 0.005 were selected.

### Subcellular fractionation

HEK293, HCT116, MDA-MB-231 or U2OS cells cultured in 6-well plates were lysed using ProteoExtract Subcellular Proteome Extraction Kit (Sigma, #539790) following manufacturer instructions to obtain cytoplasmic, membrane and nuclear fractions, and used for immunoblotting analysis of the subcellular localization of AEP in the nuclear and membrane fractions, and also to analyze enzymatic activity of AEP in the nuclear fraction, as previously described[32].

### Cell proliferation analyses

HCT116, MDA-MB-231or U2OS (3 x 10^4^ cells) were cultured in 6-well plates and cultured for 24, 48 and 72 hours in the presence or absence of 50μM MVO. At the times indicated cells were detached by trypsinization, washed twice with ice-cold PBS, resuspended in 1ml of DMEM and counted using a hemocytometer. Each data point was counted at least 5-6 time, using their average to increase the accuracy of the data. Data represents the mean ± SEM from at least 5 independent, biological replicas.

### Generation of stable AEP knock-down cancer cell lines by lentiviral infection

To generate stable AEP knock-down cancer cell lines (MDA-MB-231 and U2OS), MISSION shRNA against AEP (Sigma, #SHC001) in pLKO.1-Puro (RRID:Addgene_139470) was purchased from Sigma (5’-GTATTGAGAAGGGTCATATTT-3’, #TCRN0000276301). Lentivirus were produced upon transfection of HEK293 cells using the calcium phosphate protocol using psPAX2 (lentiviral packaging plasmid) and pMD2.G (envelope expressing plasmid) (RRID:Addgene_12259) vectors. Supernatant containing the lentivirus were collected 48 h after transfection and concentrated by ultracentrifugation at 22,000 rpm for 90 min at +4°C (Beckman Coulter Optima TM L-100K). MDA-MB-231 and U2OS were plated (6 x 10^5^ cells) in 10 cm dishes and transduced with lentiviral particles carrying shRNA against AEP or scramble shRNA (5’-CTTTGGGTGATCTACGTTA-3’) 2 days later. Stable populations of stable AEP knock-down cell lines for both MDA-MB-231 and U2OS cells were obtained after selection of infected cells in high-glucose DMEM containing 1 μg/ml puromycin (Gibco, #A1113803).

### Cell cycle analyses

Stable AEP knock-down MDA-MB-231 and U2OS cells were trypsinized, washed three times using ice-cold PBS, and 2 x 10^6^ cells were fixed using 70% ethanol at - 20°C overnight. Then, cells were collected by centrifugation at 200 g for 10 min at +4°C, washed carefully three times with ice-cold PBS and then stained in 500μl of propidium iodide (PI) staining solution (0.1% Triton X-100, 20μg/ml PI (Sigma, #P4864) and 100μg/ml DNAse-free RNAse A (Sigma, #10109142001) in PBS) with incubation at 37°C for 15 min in the dark. Samples were analyzed in a LSRFortessa X-20 (BD Biosciences) ant the quantitative analyses were carried out using FlowJo software (Becton Dickinson, v.10.7.2) (RRID:SCR_008520).

### DR-GFP, SA-GFP and EJ5-GFP assays

U2OS cells stably carrying a single copy of the DR-GFP, SA-GFP or EJ5-GFP reporters were used to analyze the role of AEP in the different double-strand break repair pathways, as previously described[75]. Cells were seeded in 6-well plates and infected with lentiviral particles containing the AEP shRNA-pLKO.1 construct using 8μg/mL polybrene. The next day cells were infected with lentiviral particles harboring the I-SceI-BFP expression construct. Forty-eight hours later, cells were collected by trypsinization, fixed with 4% paraformaldehyde for 20 min and analyzed with a BD FACS LSRFortessa X-20 and the FACS DIVA software. The number of GFP-positive cells from at least 10,000 events positive for blue fluorescence (infected with the I-SceI-BFP construct) was scored. The data represents the mean and SD from at least three independent experiments.

### Kaplan-Meier survival curves and AEP expression levels in cancer

The expression levels of AEP in human solid tumors vs normal tissue samples were analyzed using the GEPIA2 online tool and The Cancer Genome Atlas (TCGA) database. Kaplan-Meier survival analyses and gene expression box plots were calculated from the GEPIA2 online tool[30]. Kaplan-Meier survival analyses were performed using the GEPIA2 online tool to interrogate the effect of changes in the expression levels of AEP in the overall survival and prognosis in different human solid tumors.

### Breast cancer patient data

Data from breast cancer patients was obtained from the TCGA database through the cBioportal website. Kaplan-Meier analyses and correlation analyses were done using GraphPad Prism software 9 (GraphPad Software, San Diego, CA, USA) (RRID:SCR_002798).

### Data representation and statistics

All data analyses, representations and statistical analyses in this manuscript were performed using GraphPad 9 (GraphPad Software, San Diego, CA, USA). Statistical significance was calculated using unpaired Student’s t-test, unless otherwise stated, using GraphPad 9.

## Data availability

All data generated in this work is available from the authors upon request. Proteomics data generated in this study are publicly available in ProteomeXchange at PXD053016. Other data generated in this study are available within the article and its supplementary data files.

## Author’s contribution

J.M-F. conceived the project, designed research, performed and analyzed experiments, wrote the manuscript, and generated funding. M.M.-H. performed and analyzed experiments and wrote the manuscript. R.V.D. wrote the manuscript and generated funding. P.H. wrote the manuscript, designed research, and generated funding. C.W. wrote the manuscript and generated funding. M.T. performed and analyzed experiments and wrote the manuscript. G.M.B. wrote the manuscript and generated funding.

## Acknowledgements

We are grateful to Kenny Milne, Douglas Lamont and the FingerPrints Proteomics Facility at the University of Dundee for mass spectrometry. We thank Dr. Reyes Sanles-Falagán for critical reading of the manuscript. Human tissue samples were obtained with the support of the MD Anderson Foundation Biobank (record number B.0000745, ISCIII National Biobank Record). J.M-F. was supported by the European Union’s Horizon 2020 research and innovation programme under the Marie Sklodowska-Curie programme (101025429), the Ministerio de Ciencia, Innovación y Universidades, Agencia Estatal de Investigación (MICINN-AEI, RYC2021-032389), the Consejería de Transformación Económica, Industria, Conocimiento y Universidades under the EMERGIA programme (EMC21_00124) from the Andalusian Regional Government and the VII PPIT of the University of Seville (2023/00000479). RVD and G.M-B were supported by MICINN-AEI and European Regional Development Fund (project PID2021-124251OB-I00 and PID2022-136854OB-I00, respectively). G.M-B was also funded by The Instituto de Salud Carlos III IISCIII, CIBERONC (CB16/12/00295) and ERA PerMed ERA-NET co-funded by the NextGeneration-EU (ISCIII and Fundacion cientifica AECC, AC21_2/00020). PH was supported by R+D+I grant (PID2019-104195G) from the Spanish Ministry of Science and Innovation-Agencia Estatal de Investigación/10.13039/501100011033. CW was supported by a Welcome Trust Programme Grant.

---

**Supplementary Figure 1.**
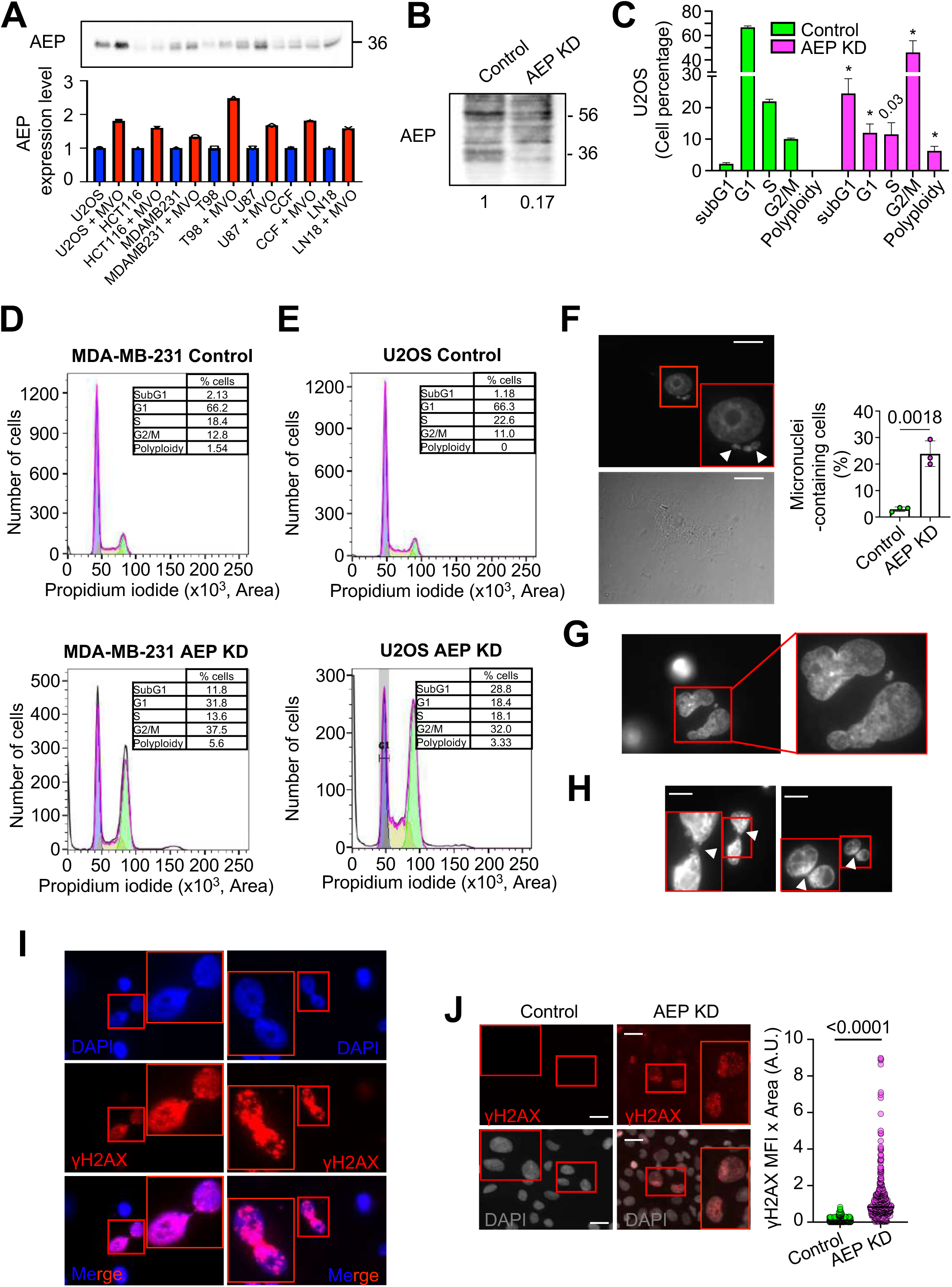
AEP deficiency in cancer cells leads to cell cycle arrest and cell death. **(A)** Immunoblot of AEP in different cancer cell lines untreated or treated with 50 μM MVO for 24 h, alongside quantitation. **(B)** Immunoblot showing the shRNA-mediated AEP knock-down in U2OS cells. **(C)** Cell cycle analyses of control and AEP shRNA transduced U2OS cells. Data represents average ± SD of 4 independent, biological replicas. * *p* value <0.01. **(D)** Representative image of the cell cycle profile of control (upper panel) compared to AEP KD (lower panel) MDA-MB-231 cells indicating the percentage of cells in the different phases of the cell cycle. **(E)** Representative image of the cell cycle profile of control (upper panel) compared to AEP KD (lower panel) U2OS cells indicating the percentage of cells in the different phases of the cell cycle. **(F)** Micrographs showing the presence of micronuclei in U2OS cells upon shRNA-mediated AEP KD alongside quantitation. Quantitation represents the average ± SD of three independent experiments, each one including more than 100 cell. Size bar = 27um. **(G)** Micrographs showing examples of polyploid cells in shRNA-mediated AEP KD U2OS cells. **(H)** Micrographs showing internuclear DNA bridges in shRNA-mediated AEP KD U2OS cells. Arrowheads indicate DNA bridges. Size bar = 27um. **(I)** Micrographs showing γH2AX-positive internuclear DNA bridges in shRNA-mediated AEP KD MDA-MD-231 cells. **(J)** Micrographs showing anti-γH2AX staining in both control (left panels) and AEP shRNA transduced (right panels) U2OS cells alongside quantitation representing the median. (n>400 cells). Size bar = 27um.

**Supplementary Figure 2.**
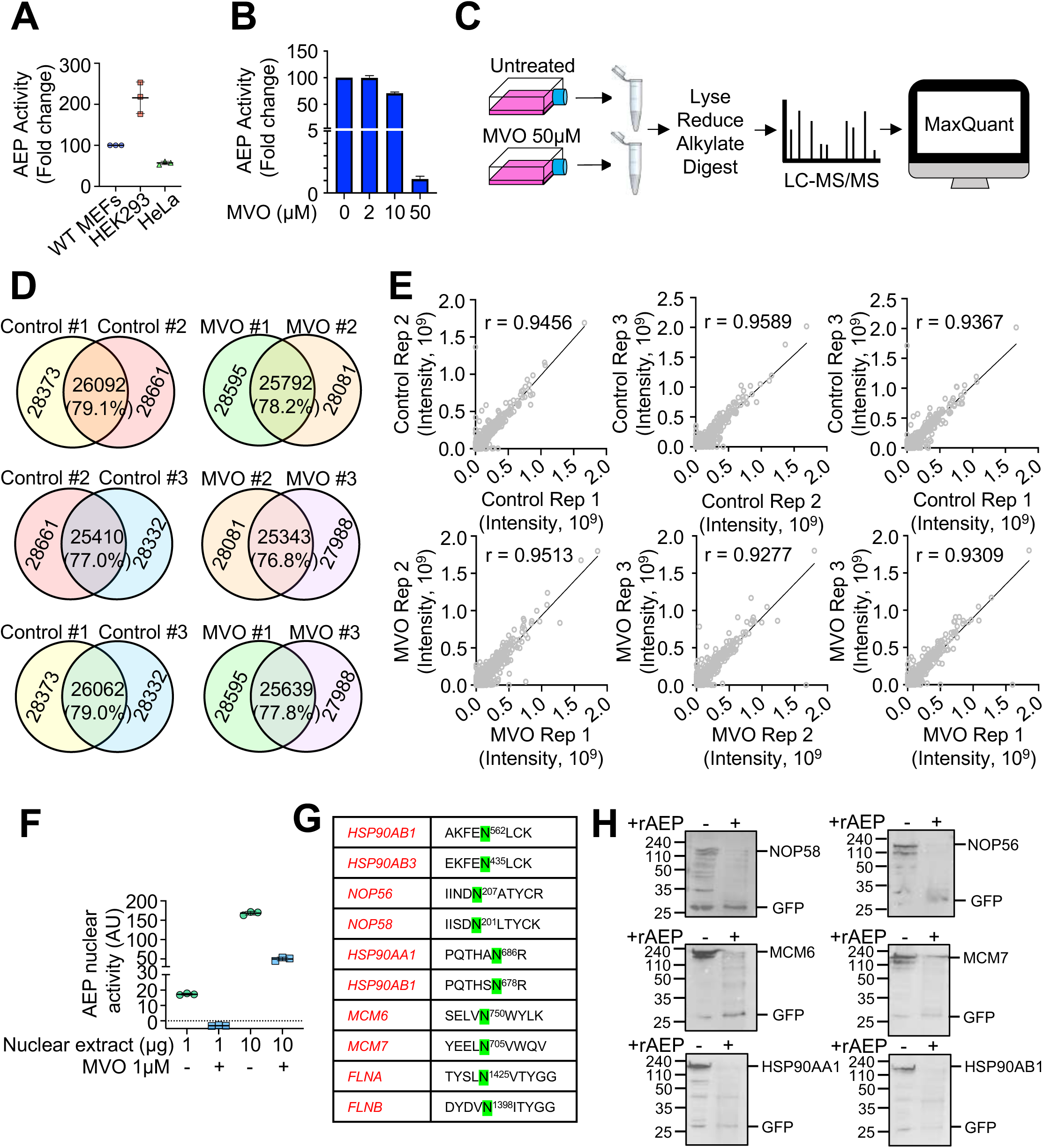
Overlap and correlation of the intensities of the peptides identified in the individual biological replicas. **(A)** AEP activity in MEFs, HEK293 and HeLa cells. **(B)** Inhibition of AEP activity in HEK293T using different concentrations of MVO26630 (0, 2, 10 or 50μM) for 16 hours. Data represents the mean ± SD of three independent, biological replicas. **(C)** Proteomics workflow. **(D)** Percentage of shared peptides identified in the individual biological replicas. (E) Correlation analyses of the intensities of the peptides identified in the individual biological replicas. **(F)** AEP activity measured in nuclear extracts of HEK293 cells and the effect of the specific AEP inhibitor (MVO26630). **(G)** Examples of Asn/Asp containing peptides accumulating upon MVO-mediated AEP inhibition highlighting the newly identified putative cleavage sites (in green). **(H)** In vitro digestion of the GFP-tagged, AEP putative targets using rAEP in pH6.5 for 1 hour.

**Supplementary Figure 3.**
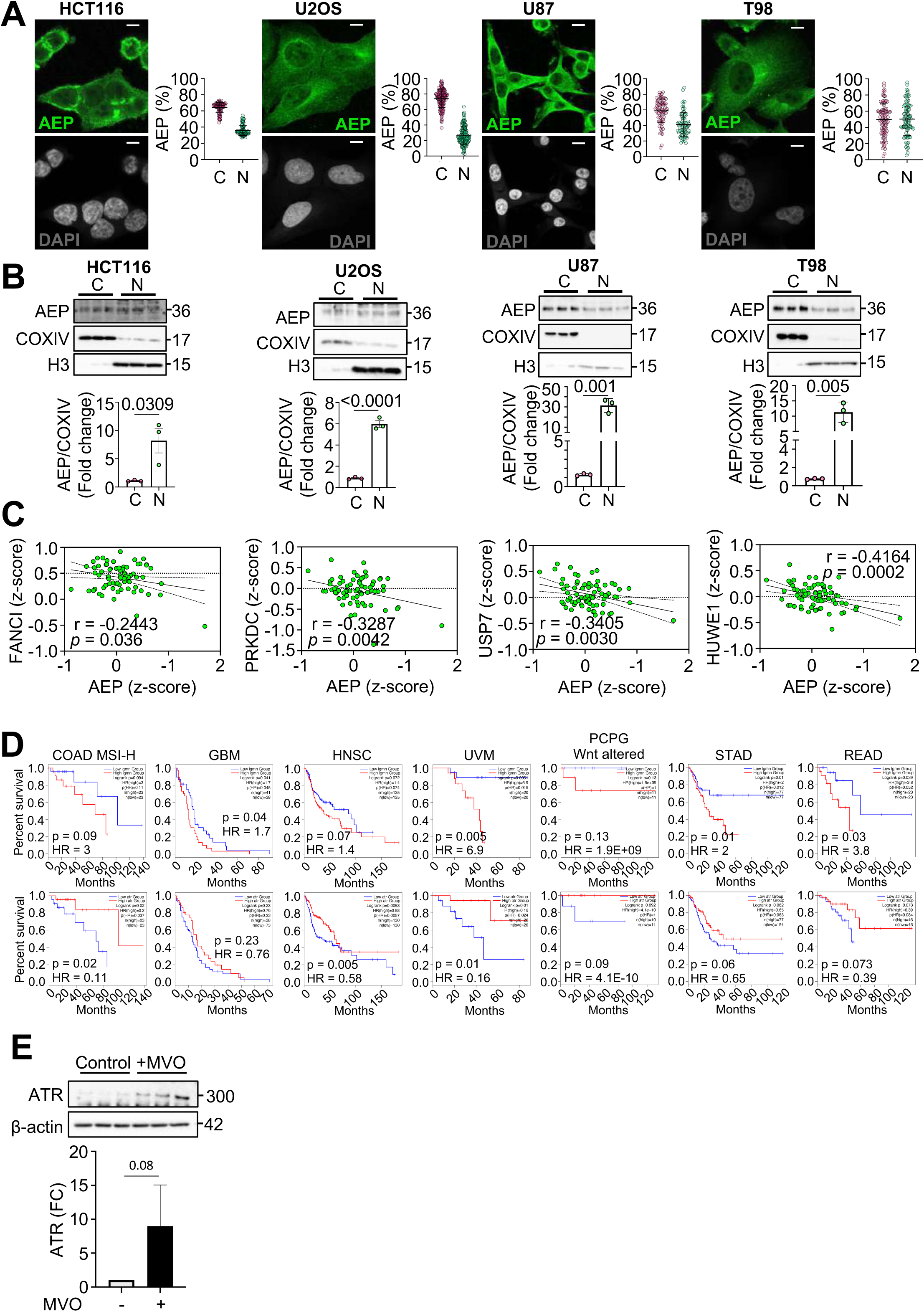
AEP regulates the levels of some of the identified targets in human cancer cells. **(A)** Immunofluorescence showing the subcellular localisation of AEP in HCT116 (n>90 cells), U2OS (n>160 cells), U87 (n>90 cells) and T98 (n>90 cells) alongside quantitation showing for each cell line the cytoplasmic (C) and nuclear (N) intensity as percentage. (Size bar=6um). **(B)** Immunoblot showing the nuclear and membrane localisation of AEP in HCT116, U2OS, U87 and T98 cells, alongside quantitation. Data represents the average ± SD of at least three independent, biological replicas. **(C)** Correlation analysis of the protein expression levels of some of the novel AEP targets identified in our proteomic analysis vs AEP in BC patients using data from the TCGA database. **(D)** Kaplan-Meier analyses of patients of different types of cancer (Colon adenocarcinoma with high microsatellite instability (COAD MSI-H), Glioblastoma (GBM), Head and Neck Squamous cell Carcinoma (HNSC), Uveal Melanoma (UVM), Pheochromocytoma and Paranglioma with Wnt-altered (PCPG), Stomach Adenocarcinoma (STAD) and Rectum Adenocarcinoma (READ)) expressing high (red line) or low (blue line) levels of AEP (upper panels) or ATR (bottom panels). **(E)** Immunoblot showing the effect of MVO-mediated AEP inhibition in the levels of ATR in U2OS cells alongside quantitation. Data represents the average ± SD of three independent, biological replicas.

**Supplementary Figure 4.**
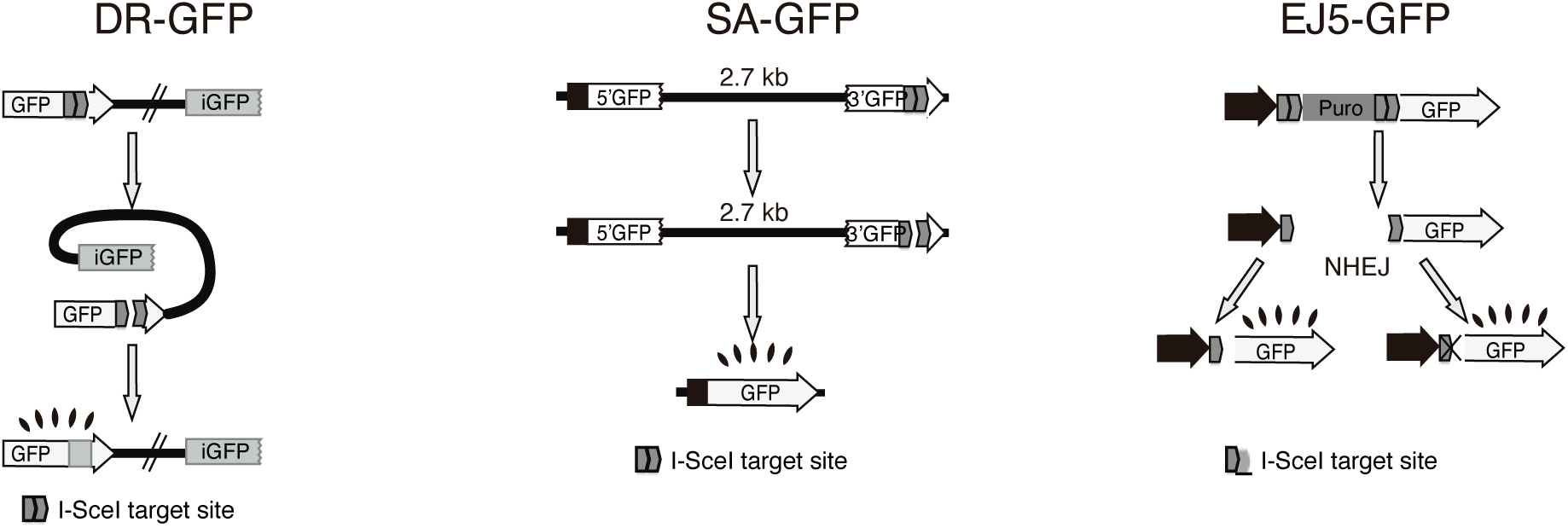
Schematic representation of the systems used to measure the efficiency of DNA repair by different DNA repair mechanisms: classical homologous recombination (DR-GFP), single-strand annealing (SA-GFP) and non-homologous end-joining (EJ5-GFP).

**Supplementary Figure 5.**
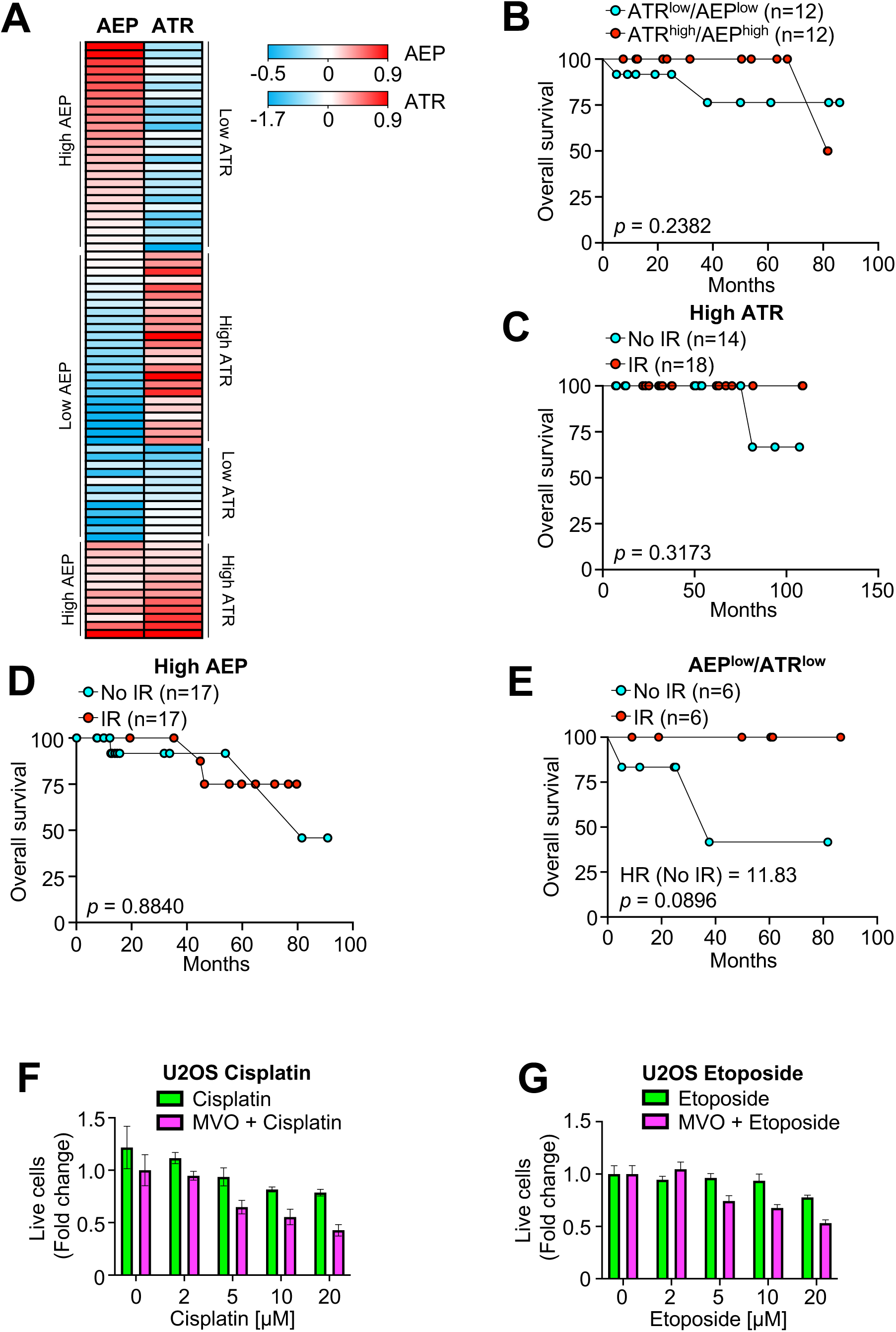
AEP inhibition sensitizes cancer cells to genotoxic stress. **(A)** Heatmap showing the levels of protein (AEP and ATR) expressed in breast cancer patients dividing patients in four groups (AEP^high^/ATR^low^; AEP^low^/ATR^high^; AEP^low^/ATR^low^; AEP^high^/ATR^high^). **(B)** Kaplan-Meier analysis in AEP^low^/ATR^low^ (cyan dots) or AEP^high^/ATR^high^ (red dots) breast cancer patients at the protein level. **(C)** Kaplan-Meier analysis in breast cancer patients expressing high ATR protein levels treated (red dots) or untreated (cyan dots) with radiation. **(D)** Kaplan-Meier analysis in breast cancer patients expressing high protein AEP levels treated (red dots) or untreated (cyan dots) with radiation. **(E**) Kaplan-Meier analysis in AEP^low^/ATR^low^ breast cancer patients treated (red dots) or untreated (cyan dots) with radiation. **(F)** Dose response of cisplatin in U2OS cell in the presence (magenta bars) or absence of MVO (green bars). **(G**) Dose response of etoposide in U2OS cell in the presence (magenta bars) or absence of MVO (green bars).

**Supplementary Figure 6.**
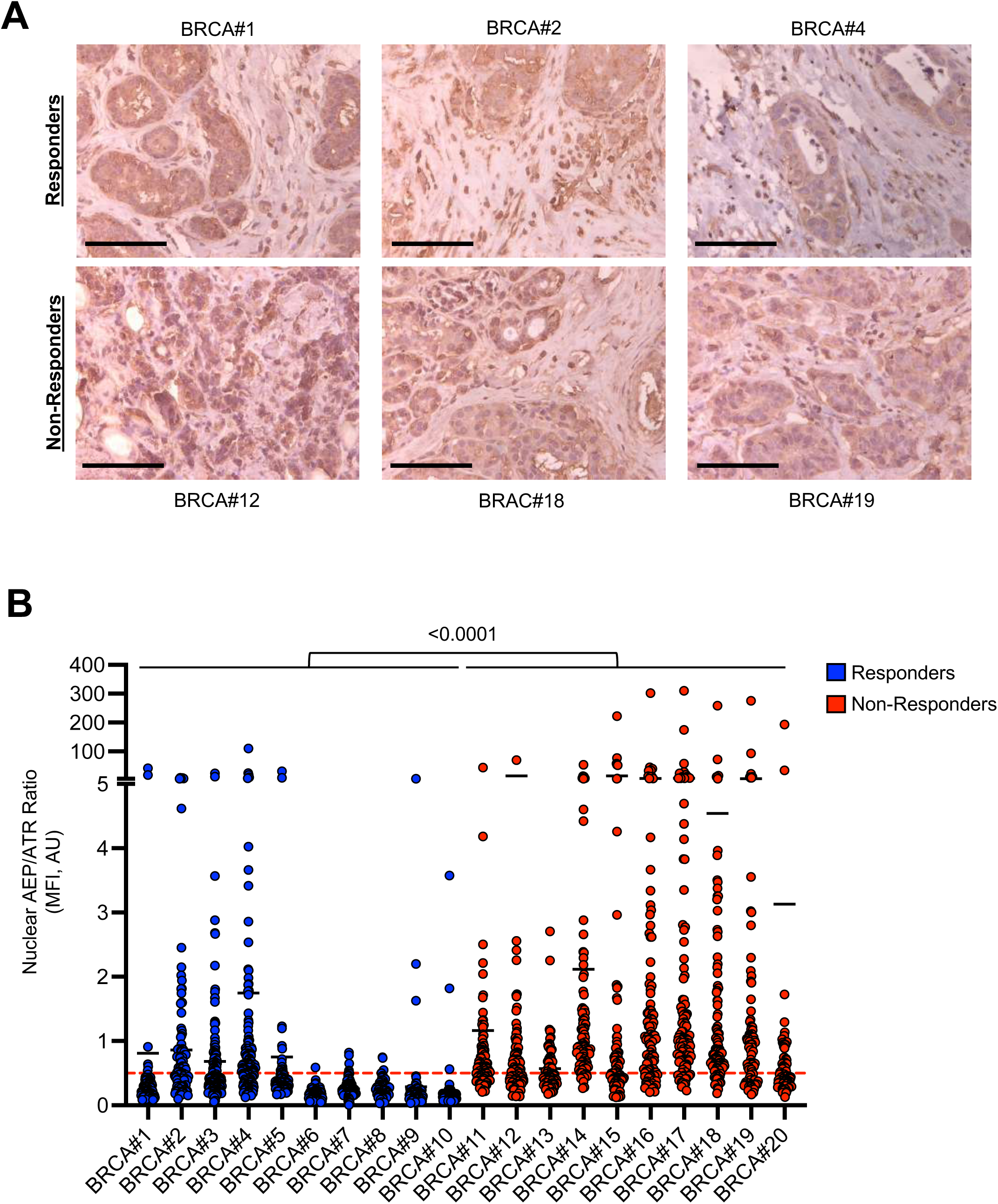
Increased AEP/ATR levels in ductal invasive breast carcinoma non-responder patients. **(A)** Microscopy images (40x) of the immunohistochemical analysis of AEP expression in responder and non-responder invasive ductal breast carcinoma samples (Size bar = 40μm). **(B)** Nuclear AEP/ATR ratio obtained from responder (n=10, blue dots) and non-responder (n=10, red dots) invasive ductal breast carcinoma patients. Each dot represents the nuclear AEP/ATR ratio obtained for each ductal breast carcinoma cell (n>80 cells per patient). Two-way ANOVA was used to calculate the statistical significance between responder and non-responder groups. Red dotted line indicates a nuclear AEP/ATR ratio greater than 0.5.

